## Supplementary data for "Sequence-sensitive elastic network captures dynamical features necessary for miR-125a maturation"

October 28, 2022

Contact:

### 1 SUPPLEMENTARY DATA

#### 1.1 *Molecular dynamics simulations*

We performed molecular dynamics simulations for the 16 possible miR-125a variants at base pair 22, the position of the G22U population SNP leading to poor prognisis in breast cancer [1]. We used the Chen-Garcia forcefield [2] with the GROMACS software [3] to accumulate more than 100ns for three replicate trajectories per variant, for a total of 5.75$\mu$s simulation data. Supplementary Table 6 gives the simulation time of every individual trajectory. The starting structures were generated from MC-Sym model 22 because previous analyses had shown this model to lead to statistically significant performance in predicting maturation efficiency from the Dynamical Signature distance. This analysis, as well as the MD simulations, were performed before we developed the novel approach of using the full Dynamical Signature inside LASSO regression.

Starting from the *in silico* structures mutated using ModeRNA [4], we used a standard cubic box with edges 1nm away from the miR-125a structure, solvated the RNA using TIP4P water following personal communication with Alan Chen. 100mM NaCL was added to the system, in addition to the necessary amount of cations to make the system neutral. Energy minimization was performed until the biggest force on any atom was under 1 kJ/mol, followed by NVT and NPT equilibration for 100ps each, at 310K (human body temperature). The simulation was then carried until at least 100 ns of simulation time had been acquired, also at 310K, with a 2fs time-step. The hardware used was the Intel Gold 6148 Skylake 20-core CPU @ 2.4GHz, as part of Compute Canada's Béluga computing cluster. The total computational cost for the 5.75$\mu$s of simulation time was around 40 core-years.

From the MD trajectories, we computed Dynamical Signatures by taking the mean-square fluctuations at the three atomic positions on which beads are placed for the ENMs in the present study: the P, C1′ and C2 atoms. This filtering ensures better comparison between the MD trajectories and ENCoM Dynamical Signatures, in addition to providing a flexibility measure for each of the phosphate, sugar and base groups, aruguably the logical subdivision of a nucleotide into more or less rigid groups.

Because only 16 variants were simulated, training linear regression models on the full Dynamical Signatures would lead to severe overfitting since there are 257 positions in the signature. However, to see if the flexibility profiles from the MD simulations better capture dynamical patterns necessary for miR-125a maturation, we plotted the measured maturation efficiency against the euclidean distance between the WT dynamical signature and every mutant signature. Since there are three replicate trajectories per variant, we take the mean euclidean distance from the 9 pairwise distances (3 mutants X 3 WT).

Supplementary Figure 1 reports the correlation between the distance to the WT signature and the measured maturation efficiency. The expected correlation is negative: the further the dynamical profile is from the WT, the lower the maturation efficiency, giving rise to a negative correlation, as reported by Dallaire and coworkers [5]. Surprisingly, the MD signature leads to a very weak positive correlation. While the ENCoM EntroSigs do not reach statistical significance at R=-0.27, the correlation is of the expected sign. None of the four correlations reported reach statistical significance, however ENCoM EntroSigs perform better than the MD MSF.

We thus conclude that at the simulated timescales, MD simulations do not provide useful information over the ENCoM conformational space. Perhaps longer timescales would lead to better performance, however let us remind that the goal of our novel approach is the high-throughput computational prediction of variant effects. Since the computational cost of one ENCoM Entropic Signature is around 3 seconds on a single CPU, it represents a speedup of $8.8 * 10^6$ over the MD approach at the timescales presented here, namely slightly above 100ns per trajectory. The MD trajectories, converted to PDB format with the medoid frame from each 1ns window, are available online.

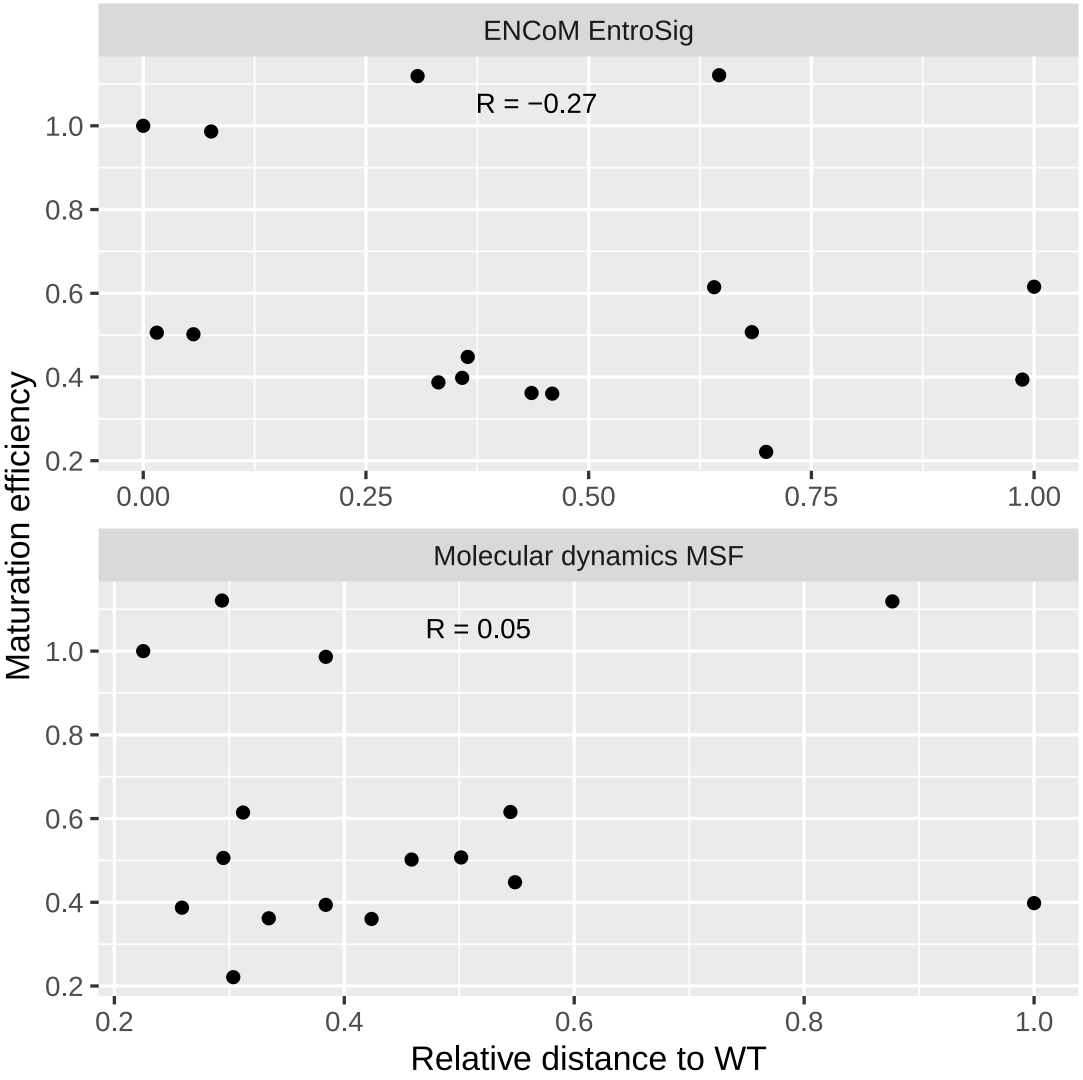


**Supplementary Figure 1: Correlation between miR-125a maturation efficiency and Dynamical Signature distance to WT for both ENCoM and MD simulations.** The average distance for all 9 possibilities is reported for the MD simulations (3 replicates for the variant, 3 replicates for WT). The mean-square fluctuation was computed at P, C1′ and C2 atoms as the square of the RMSF computed with GROMACS. The ENCoM Entropic Signature was computed at $\beta = e^{0.25}$, the value selected from the B-factors benchmark. The Pearson linear correlation coefficients reported do not reach statistical significance.

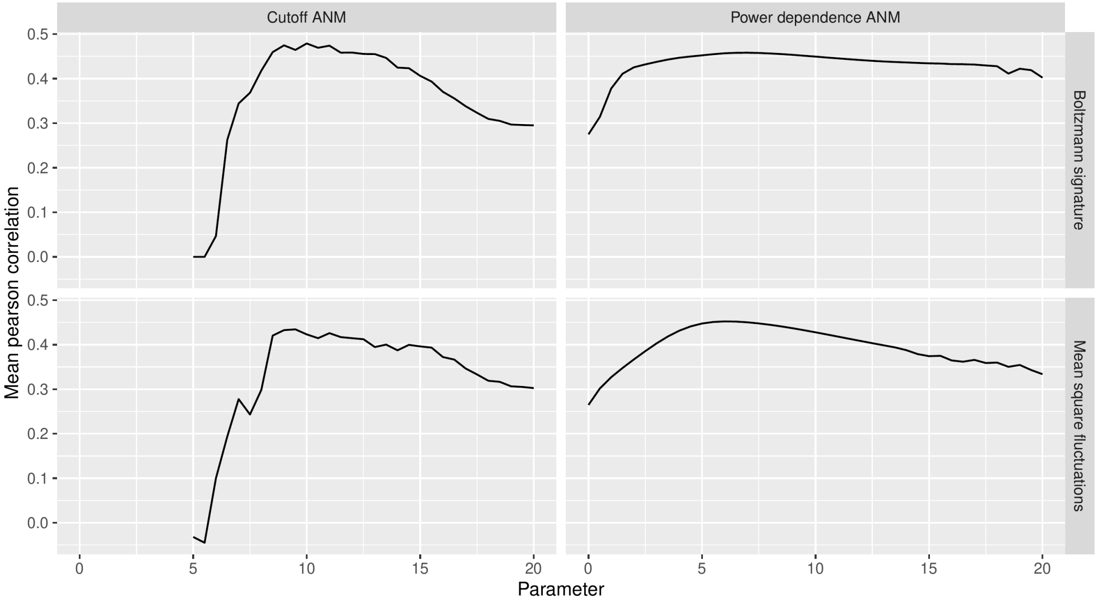


**Supplementary Figure 2: Cut-ANM and PD-ANM parameter sweep for the B-factors benchmark.** Parameters are interaction distance cutoff (in Å) for Cut-ANM and power dependence for PD-ANM. Both are varied in increments of 0.5 from their minimum practical value up to 20.

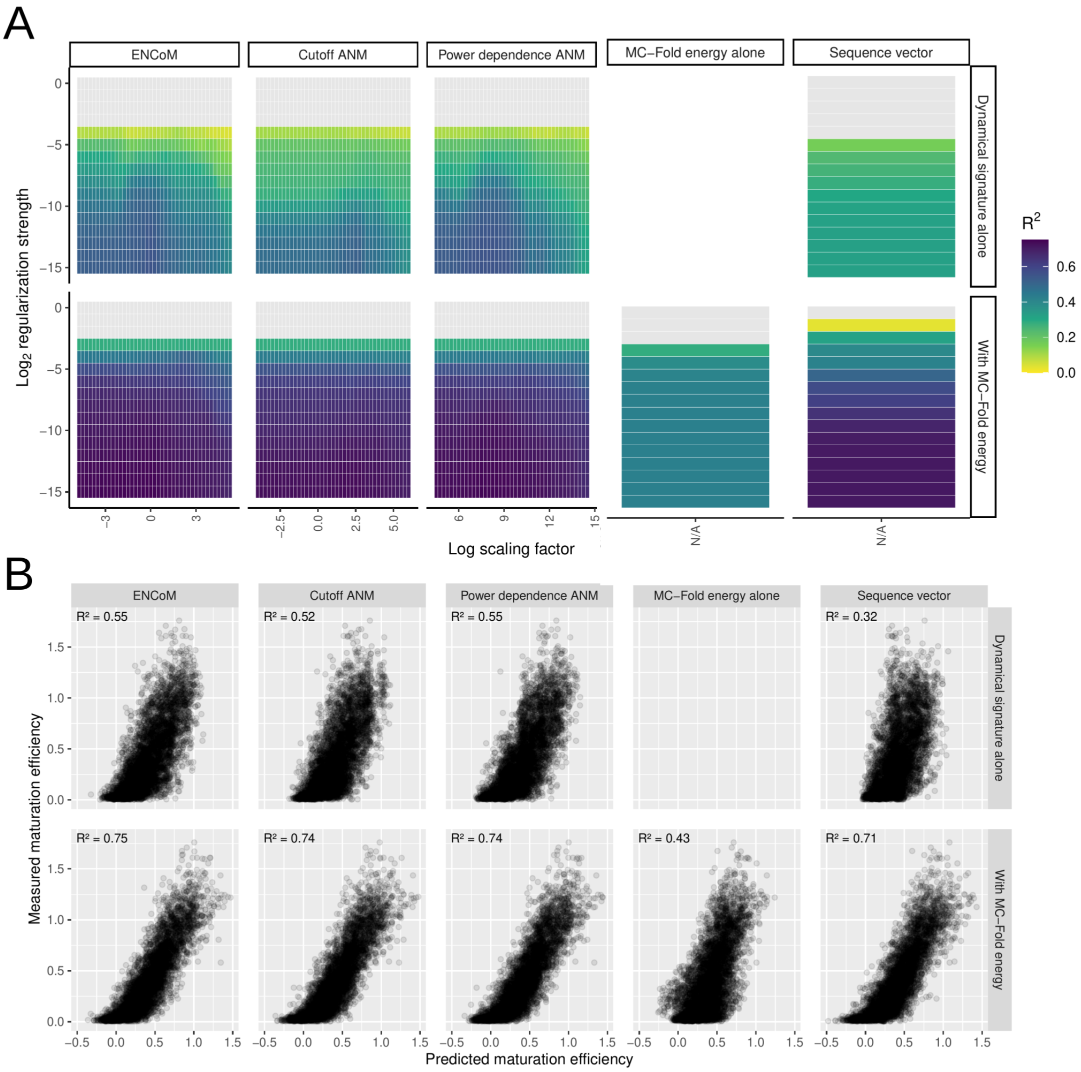


**Supplementary Figure 3: Performance of multiple linear regression on an 80-20 train-test split.** A) Predictive R2 for each model alone or in combination with the MC-Fold enthalpy of folding. For the three ENMs, scaling factors for the dynamical signature were explored around the value which gave the best respective performance in the B-factors benchmark. Predictive R2 values below 0 are shown in gray. B) Performance on the test set for the best combination of parameters for every model alone or in tandem with MC-Fold.

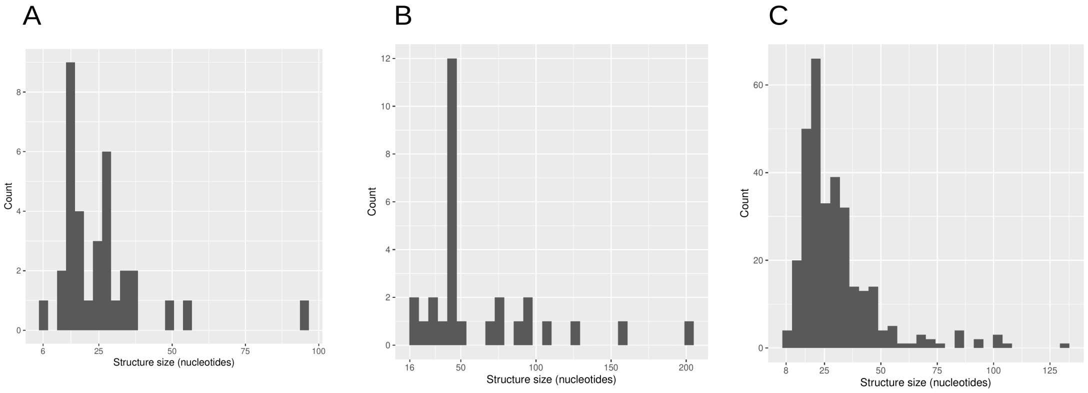


**Supplementary Figure 4: Size distributions of the sequence clusters from the 3 benchmarks.** A) B-factors benchmark. B) Conformational change from X-ray structures benchmark. C) NMR ensemble structural variance benchmark.

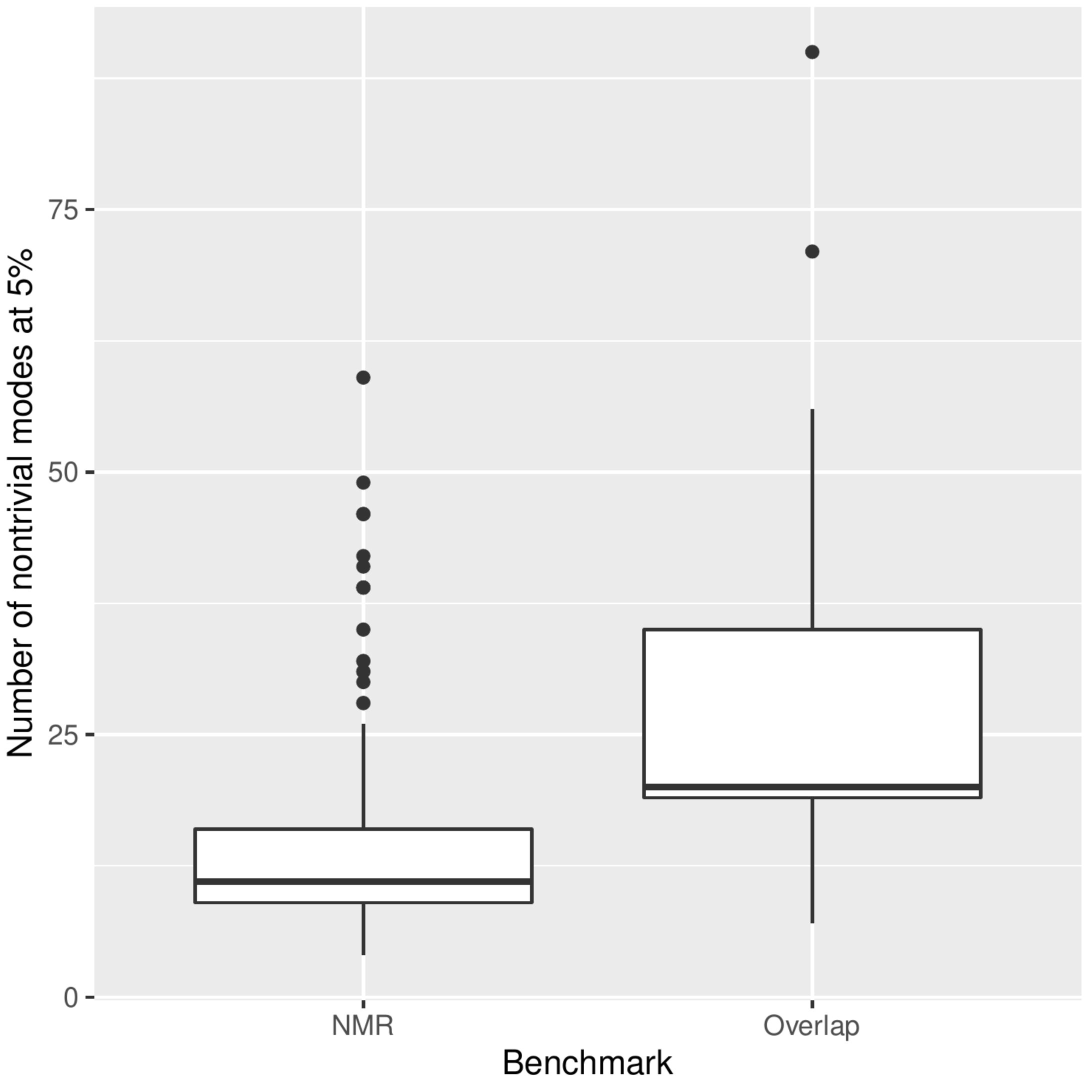


**Supplementary Figure 5: Number of nontrivial normal modes at 5% for the overlap and NMR benchmarks.**

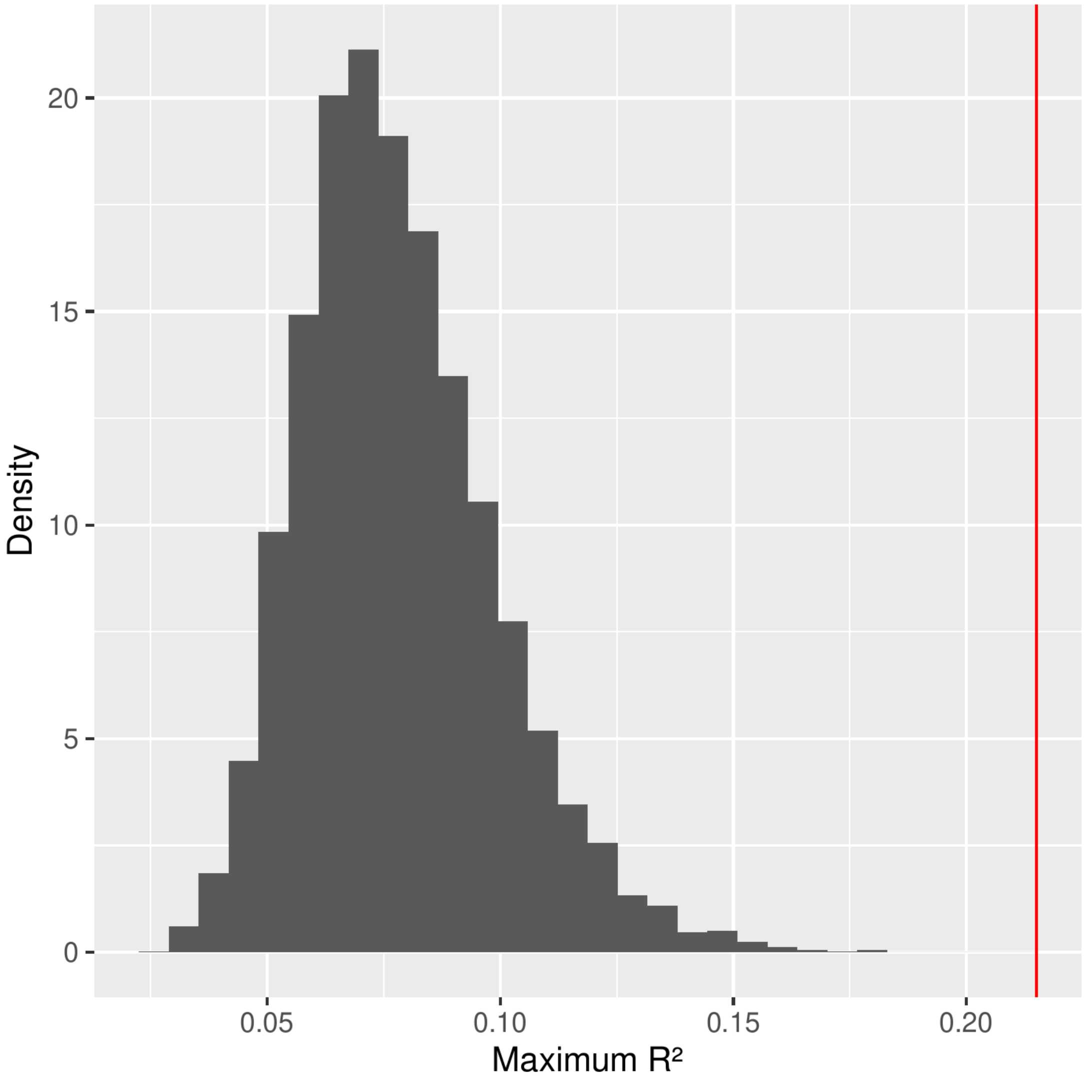


**Supplementary Figure 6: Simulation of predictive R-squared from combinations of MC-Fold enthalpy with random noise.** Each of the 10 000 iterations represents the best predictive R2 value from 656 samplings of 116 values from a gaussian distribution centered around the training set mean and with a standard deviation two times smaller. The noise is then averaged at each sampling with the MC-Fold predictions and predictive R2 is computed. The red line shows the R2 of the ENCoM-MC-Fold combination on the hard benchmark (0.215).

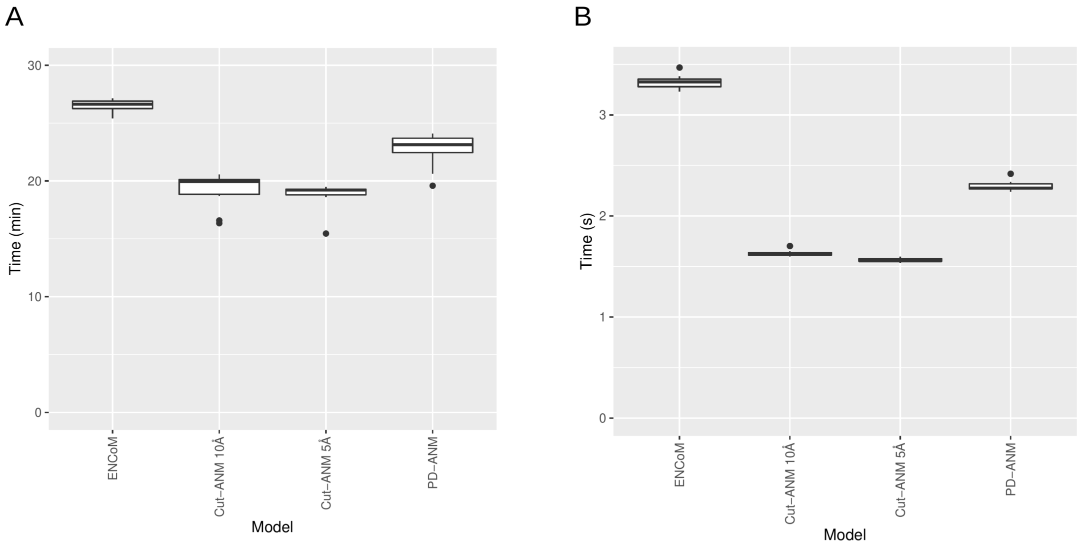


**Supplementary Figure 7: Computational cost of the ENMs on the Thermus thermophilus 30S ribosomal subunit and on miR-125a.** A) Thermus thermophilus 30S ribosomal subunit. B) MiR-125a.

**Supplementary Table 1:** Atom type assignation of the four standard nucleotides. The atom types assigned are the ones from Sobolev and coworkers, divided in eight classes: hydrophilic, acceptor, donor, hydrophobic, aromatic, neutral, neutral-donor and neutral-acceptor.

| Atom name | Atom type |
|---|---|
| Nucleobase atoms | |
| C2-C6, C8 | Aromatic |
| N2, N4, N6 | Donor |
| N3, N7 | Acceptor |
| N9 | Neutral |
| Sugar atoms | |
| C1′, C3′-C5′ | Neutral-acceptor |
| C2′ | Neutral |
| O4′ | Acceptor |
| O2′ | Hydrophilic |
| Phosphate atoms | |
| P | Neutral-acceptor |
| OP1-OP3, O5′, O3′ | Acceptor |
| Specific atom types | |
| Adenine | |
| N1 | Acceptor |
| Guanine | |
| N1 | Donor |
| Cytidine | |
| N1 | Neutral |
| N5 | Acceptor |
| O2 | Acceptor |
| Uridine | |
| O2 | Acceptor |
| O4 | Acceptor |

**Supplementary Table 2:** PDB codes of the structures used for the three benchmarks.

| | | | | | | | | | |
|---|---|---|---|---|---|---|---|---|---|
| B-factors prediction | | | | | | | | | |
| 157D | 1CSL | 1DQH | 1I9X | 1KD5 | 1MSY | 1OSU | 1Q9A | 1RNA | 1RXB |
| 1ZEV | 255D | 259D | 280D | 2OE6 | 2V6W | 2V7R | 2VUQ | 2XSL | 353D |
| 3GM7 | 3SZX | 402D | 405D | 413D | 433D | 438D | 472D | 480D | 483D |
| 4C40 | 4E59 | 4U37 | 4U38 | 5C5W | 5NXT | 5VGW | 7EAG | | |
| Conformational change prediction | | | | | | | | | |
| 1HR2 | 1J7T | 1L8V | 1LC4 | 1MWL | 1NLC | 1O3Z | 1O9M | 1XP7 | 1XPE |
| 1XPF | 1Y3S | 1YRJ | 1YXP | 1ZCI | 1ZX7 | 1ZZ5 | 2A04 | 2B8R | 2B8S |
| 2BE0 | 2BEE | 2ESJ | 2ET3 | 2ET4 | 2ET5 | 2ET8 | 2F4S | 2F4T | 2F4U |
| 2FCX | 2FCY | 2FCZ | 2FD0 | 2FQN | 2G5K | 2GPM | 2NOK | 2O3W | 2O3X |
| 2OIJ | 2OIY | 2OJ0 | 2PN4 | 2PWT | 2QEK | 3BNL | 3BNQ | 3BNR | 3BNS |
| 3C44 | 3CW5 | 3CW6 | 3DVV | 3FAR | 3GCA | 3L0U | 3OWI | 3OWW | 3OWZ |
| 3OX0 | 3OXB | 3OXD | 3OXE | 3OXJ | 3OXM | 3TD0 | 3TD1 | 3WRU | 439D |
| 462D | 4F8U | 4F8V | 4GPY | 4K31 | 4K32 | 4L81 | 4MSB | 4MSR | 4OQU |
| 4P20 | 4P3S | 4P3T | 4P3U | 4P43 | 4PDQ | 4RZD | 4TZX | 4TZY | 4XNR |
| 4Y1J | 5E54 | 5L4O | 5SWD | 5SWE | 5TDK | 5UZA | 5XZ1 | 5ZEG | 5ZEI |
| 5ZEJ | 5ZEM | 6BJX | 6C8D | 6CAB | 6CB3 | 6D8L | 6D8M | 6D8O | 6DLR |
| 6DLS | 6DNR | 6E80 | 6E81 | 6E82 | 6E84 | 6JBG | 6N5K | 6N5O | 6VUH |
| 6VWT | 6VWV | 6WY3 | 6XKN | 6XKO | 6Y3G | 6Z18 | 7EOI | 7EOK | |
| NMR ensemble variance prediction | | | | | | | | | |
| 17RA | 1A3M | 1A51 | 1A60 | 1A9L | 1AFX | 1ANR | 1AQO | 1ATO | 1ATV |
| 1ATW | 1B36 | 1BGZ | 1BN0 | 1BVJ | 1CQL | 1D0U | 1E4P | 1E95 | 1EBQ |
| 1EBR | 1EBS | 1ELH | 1ESY | 1F5G | 1F5H | 1F7F | 1F7G | 1F84 | 1F85 |
| 1FHK | 1FYO | 1GUC | 1HLX | 1HWQ | 1I3X | 1IDV | 1IE1 | 1IE2 | 1IKD |
| 1JO7 | 1JOX | 1JP0 | 1JTJ | 1JU1 | 1JU7 | 1JUR | 1K4A | 1K4B | 1K5I |
| 1K6G | 1K6H | 1KKA | 1KKS | 1LC6 | 1LDZ | 1LMV | 1M5L | 1M82 | 1MFJ |
| 1MFK | 1MFY | 1MNX | 1MT4 | 1N66 | 1N8X | 1NA2 | 1NBR | 1NC0 | 1OQ0 |
| 1OSW | 1OW9 | 1P5M | 1P5N | 1P5O | 1PJY | 1Q75 | 1QC8 | 1QES | 1QET |
| 1QWA | 1QWB | 1R2P | 1R7W | 1R7Z | 1RNG | 1ROQ | 1RRR | 1S34 | 1S9S |
| 1SCL | 1SLP | 1SY4 | 1SYZ | 1T4X | 1TBK | 1TJZ | 1TLR | 1TXS | 1U3K |
| 1UUU | 1VOP | 1WKS | 1XHP | 1YLG | 1YMO | 1YN1 | 1YNC | 1YNE | 1YNG |
| 1YSV | 1Z2J | 1Z30 | 1Z31 | 1ZC5 | 1ZIF | 1ZIG | 1ZIH | 28SP | 2ADT |
| 2AHT | 2B7G | 2D17 | 2D18 | 2D19 | 2ES5 | 2EUY | 2EVY | 2F4X | 2F87 |
| 2F88 | 2FDT | 2FEY | 2G1W | 2GIO | 2GIP | 2GM0 | 2GRW | 2GV3 | 2GV4 |
| 2GVO | 2H49 | 2HEM | 2HNS | 2HUA | 2IRN | 2IRO | 2IXY | 2IXZ | 2JR4 |
| 2JSE | 2JTP | 2JWV | 2JXQ | 2JXS | 2JXV | 2JYF | 2JYH | 2JYJ | 2JYM |
| 2K3Z | 2K41 | 2K5Z | 2K65 | 2K66 | 2K95 | 2K96 | 2KBP | 2KD8 | 2KE6 |
| 2KEZ | 2KF0 | 2KHY | 2KOC | 2KPC | 2KPD | 2KPV | 2KRL | 2KRP | 2KRQ |
| 2KUR | 2KUU | 2KUW | 2KVN | 2KXZ | 2KY0 | 2KY1 | 2KY2 | 2KYD | 2KZL |
| 2L1F | 2L2J | 2L3E | 2L5Z | 2L6I | 2L8F | 2LAC | 2LBJ | 2LBK | 2LBL |
| 2LC8 | 2LDL | 2LDT | 2LHP | 2LJJ | 2LK3 | 2LP9 | 2LPA | 2LPS | 2LPT |
| 2LQZ | 2LU0 | 2LUB | 2LUN | 2LV0 | 2LX1 | 2M12 | 2M18 | 2M21 | 2M22 |
| 2M23 | 2M24 | 2M4W | 2M57 | 2M5U | 2M8K | 2MEQ | 2MFD | 2MI0 | 2MNC |
| 2MQT | 2MTJ | 2MXJ | 2MXK | 2MXL | 2N2O | 2N2P | 2N3Q | 2N4L | 2N6S |
| 2N6T | 2N6W | 2N6X | 2N7M | 2N7X | 2N8V | 2NBX | 2NBY | 2NBZ | 2NC0 |
| 2NC1 | 2NCI | 2O33 | 2OJ7 | 2OJ8 | 2P89 | 2PCV | 2PCW | 2QH2 | 2QH3 |
| 2QH4 | 2RLU | 2RN1 | 2RPT | 2RQJ | 2RVO | 2XEB | 2Y95 | 3PHP | 4A4S |
| 4A4T | 4A4U | 5A17 | 5A18 | 5IEM | 5KH8 | 5KMZ | 5KQE | 5LSN | 5N5C |
| 5UF3 | 5UZT | 5V16 | 5V17 | 5V2R | 5VH7 | 5VH8 | 5WQ1 | 6AAS | 6BG9 |
| 6BY4 | 6BY5 | 6GE1 | 6HYK | 6K84 | 6MXQ | 6N8F | 6N8I | 6NOA | 6PK9 |
| 6U79 | 6VA1 | 6VAR | 6VZC | 6W3M | 6XWJ | 6XWW | 6XXA | 6XXB | 7DD4 |
| 7JU1 | 7K4L | 7LVA | | | | | | | |

**Supplementary Table 3:** Performance on individual structures for the B-factors correlation benchmark.

| PDB code | Cluster index | ENCoM BZ | ENCoM MSF | Cut-ANM BZ | Cut-ANM MSF | PD-ANM BZ | PD-ANM MSF |
|---|---|---|---|---|---|---|---|
| 472D | 1 | 0.49 | 0.49 | 0.55 | 0.52 | 0.55 | 0.55 |
| 4U37 | 2 | 0.78 | 0.79 | 0.71 | 0.68 | 0.65 | 0.66 |
| 433D | 3 | 0.59 | 0.60 | 0.45 | 0.57 | 0.42 | 0.50 |
| 405D | 4 | 0.52 | 0.45 | 0.60 | 0.46 | 0.61 | 0.56 |
| 1CSL | 5 | 0.40 | 0.36 | 0.32 | 0.17 | 0.28 | 0.35 |
| 1Q9A | 6 | 0.44 | 0.37 | 0.65 | 0.38 | 0.56 | 0.49 |
| 480D | 6 | 0.65 | 0.65 | 0.76 | 0.66 | 0.70 | 0.67 |
| 483D | 6 | 0.39 | 0.33 | 0.56 | 0.34 | 0.50 | 0.44 |
| 5C5W | 7 | 0.69 | 0.77 | 0.45 | 0.77 | 0.47 | 0.77 |
| 2VUQ | 8 | 0.83 | 0.80 | 0.78 | 0.72 | 0.71 | 0.73 |
| 1I9X | 9 | 0.31 | 0.31 | 0.19 | 0.07 | 0.12 | 0.08 |
| 1RNA | 10 | 0.55 | 0.51 | 0.58 | 0.41 | 0.57 | 0.52 |
| 413D | 11 | 0.68 | 0.70 | 0.66 | 0.80 | 0.68 | 0.74 |
| 280D | 12 | 0.21 | 0.06 | 0.31 | 0.26 | 0.31 | 0.05 |
| 157D | 13 | 0.25 | 0.17 | 0.56 | 0.46 | 0.54 | 0.51 |
| 3SZX | 14 | 0.35 | 0.38 | 0.28 | 0.41 | 0.29 | 0.36 |
| 353D | 15 | -0.06 | -0.09 | 0.15 | -0.05 | 0.12 | 0.12 |
| 255D | 16 | 0.68 | 0.64 | 0.70 | 0.77 | 0.71 | 0.71 |
| 1ZEV | 17 | 0.31 | 0.25 | 0.10 | -0.05 | 0.04 | 0.01 |
| 3GM7 | 17 | 0.43 | 0.35 | 0.31 | 0.34 | 0.32 | 0.31 |
| 1KD5 | 18 | 0.18 | 0.16 | 0.44 | 0.19 | 0.38 | 0.25 |
| 5VGW | 19 | 0.09 | 0.03 | 0.37 | 0.35 | 0.31 | 0.18 |
| 4C40 | 20 | 0.62 | 0.50 | 0.70 | 0.44 | 0.75 | 0.81 |
| 402D | 21 | 0.42 | 0.43 | 0.40 | 0.44 | 0.42 | 0.43 |
| 1RXB | 22 | 0.59 | 0.60 | 0.64 | 0.65 | 0.60 | 0.63 |
| 259D | 22 | 0.57 | 0.58 | 0.59 | 0.63 | 0.54 | 0.58 |
| 2V6W | 23 | 0.71 | 0.73 | 0.73 | 0.66 | 0.73 | 0.74 |
| 2XSL | 24 | 0.45 | 0.47 | 0.50 | 0.48 | 0.53 | 0.50 |
| 1DQH | 25 | 0.44 | 0.46 | 0.58 | 0.54 | 0.54 | 0.52 |
| 7EAG | 26 | -0.08 | -0.20 | 0.09 | -0.09 | 0.01 | -0.09 |
| 438D | 27 | 0.47 | 0.42 | 0.54 | 0.46 | 0.52 | 0.56 |
| 5NXT | 28 | -0.01 | -0.01 | 0.13 | 0.13 | 0.11 | 0.04 |
| 1MSY | 29 | 0.63 | 0.68 | 0.65 | 0.76 | 0.63 | 0.64 |
| 2V7R | 30 | 0.82 | 0.78 | 0.79 | 0.72 | 0.71 | 0.74 |
| 4U38 | 31 | 0.28 | 0.23 | 0.29 | 0.20 | 0.32 | 0.24 |
| 2OE6 | 32 | 0.48 | 0.42 | 0.44 | 0.39 | 0.41 | 0.43 |
| 1OSU | 33 | 0.20 | 0.17 | 0.23 | 0.24 | 0.32 | 0.29 |
| 4E59 | 34 | 0.58 | 0.59 | 0.64 | 0.60 | 0.58 | 0.58 |

**Supplementary Table 4:** Performance on individual pairs of conformations for the X-ray conformational change benchmark.

| Input PDB code | Target PDB code | RMSD | Cluster index | ENCoM | Cut-ANM | PD-ANM |
|---|---|---|---|---|---|---|
| 1HR2 | 1L8V | 2.26 | 1 | 0.66 | 0.54 | 0.57 |
| 1HR2 | 6BJX | 2.21 | 1 | 0.52 | 0.44 | 0.45 |
| 1HR2 | 6D8L | 2.22 | 1 | 0.52 | 0.45 | 0.46 |
| 1HR2 | 6D8M | 2.15 | 1 | 0.56 | 0.48 | 0.52 |
| 1HR2 | 6D8O | 2.01 | 1 | 0.56 | 0.46 | 0.49 |
| 1L8V | 1HR2 | 2.26 | 1 | 0.58 | 0.55 | 0.55 |
| 1L8V | 6BJX | 2.24 | 1 | 0.58 | 0.53 | 0.57 |
| 1L8V | 6D8L | 2.25 | 1 | 0.58 | 0.53 | 0.56 |
| 1L8V | 6D8M | 2.14 | 1 | 0.63 | 0.58 | 0.59 |
| 1L8V | 6D8O | 2.14 | 1 | 0.64 | 0.57 | 0.58 |
| 6BJX | 1HR2 | 2.21 | 1 | 0.50 | 0.42 | 0.45 |
| 6BJX | 1L8V | 2.24 | 1 | 0.60 | 0.51 | 0.53 |
| 6D8L | 1HR2 | 2.22 | 1 | 0.50 | 0.43 | 0.44 |
| 6D8L | 1L8V | 2.25 | 1 | 0.58 | 0.50 | 0.52 |
| 6D8M | 1HR2 | 2.15 | 1 | 0.53 | 0.46 | 0.47 |
| 6D8M | 1L8V | 2.14 | 1 | 0.63 | 0.57 | 0.57 |
| 6D8O | 1HR2 | 2.01 | 1 | 0.53 | 0.46 | 0.46 |
| 6D8O | 1L8V | 2.14 | 1 | 0.64 | 0.55 | 0.57 |
| 1J7T | 1O9M | 2.02 | 2 | 0.63 | 0.66 | 0.67 |
| 1J7T | 2ET8 | 2.02 | 2 | 0.59 | 0.57 | 0.60 |
| 1J7T | 3BNL | 2.90 | 2 | 0.77 | 0.80 | 0.81 |
| 1J7T | 4F8U | 2.36 | 2 | 0.92 | 0.90 | 0.92 |
| 1LC4 | 3BNL | 2.96 | 2 | 0.78 | 0.84 | 0.83 |
| 1LC4 | 4F8U | 2.07 | 2 | 0.91 | 0.92 | 0.92 |
| 1MWL | 1O9M | 2.14 | 2 | 0.63 | 0.66 | 0.66 |
| 1MWL | 2ET8 | 2.24 | 2 | 0.63 | 0.61 | 0.63 |
| 1MWL | 3BNL | 3.32 | 2 | 0.80 | 0.83 | 0.85 |
| 1MWL | 4F8U | 2.25 | 2 | 0.79 | 0.75 | 0.77 |
| 1O9M | 1J7T | 2.02 | 2 | 0.63 | 0.65 | 0.65 |
| 1O9M | 1MWL | 2.14 | 2 | 0.64 | 0.65 | 0.69 |
| 1O9M | 1YRJ | 2.43 | 2 | 0.68 | 0.67 | 0.69 |
| 1O9M | 2BE0 | 2.06 | 2 | 0.64 | 0.63 | 0.68 |
| 1O9M | 2BEE | 2.05 | 2 | 0.64 | 0.63 | 0.68 |
| 1O9M | 2ET3 | 2.31 | 2 | 0.70 | 0.72 | 0.74 |
| 1O9M | 2F4S | 2.64 | 2 | 0.47 | 0.57 | 0.55 |
| 1O9M | 2F4T | 2.00 | 2 | 0.32 | 0.24 | 0.22 |
| 1O9M | 2F4U | 2.65 | 2 | 0.49 | 0.58 | 0.57 |
| 1O9M | 2O3X | 2.55 | 2 | 0.43 | 0.53 | 0.52 |
| 1O9M | 2PWT | 2.18 | 2 | 0.71 | 0.73 | 0.75 |
| 1O9M | 3BNL | 2.91 | 2 | 0.72 | 0.75 | 0.76 |
| 1O9M | 4F8U | 2.59 | 2 | 0.73 | 0.70 | 0.72 |
| 1O9M | 4F8V | 2.28 | 2 | 0.67 | 0.66 | 0.68 |
| 1YRJ | 1O9M | 2.43 | 2 | 0.69 | 0.63 | 0.60 |
| 1YRJ | 2ET8 | 2.44 | 2 | 0.68 | 0.64 | 0.63 |
| 1YRJ | 2F4S | 2.51 | 2 | 0.68 | 0.63 | 0.65 |
| 1YRJ | 2F4T | 2.47 | 2 | 0.67 | 0.62 | 0.62 |
| 1YRJ | 2F4U | 2.23 | 2 | 0.62 | 0.59 | 0.60 |
| 1YRJ | 2O3X | 2.38 | 2 | 0.67 | 0.60 | 0.63 |
| 1YRJ | 2PWT | 2.04 | 2 | 0.84 | 0.76 | 0.76 |

| Supplementary Table 4 (continued) | | | | | | |
|---|---|---|---|---|---|---|
| Input PDB code | Target PDB code | RMSD | Cluster index | ENCoM | Cut-ANM | PD-ANM |
| 1YRJ | 3BNL | 3.44 | 2 | 0.79 | 0.77 | 0.77 |
| 1YRJ | 4F8U | 2.81 | 2 | 0.88 | 0.78 | 0.78 |
| 1YRJ | 4F8V | 2.02 | 2 | 0.85 | 0.80 | 0.82 |
| 2BE0 | 1O9M | 2.06 | 2 | 0.66 | 0.65 | 0.67 |
| 2BE0 | 2ET8 | 2.23 | 2 | 0.70 | 0.64 | 0.67 |
| 2BE0 | 2F4S | 2.33 | 2 | 0.71 | 0.69 | 0.71 |
| 2BE0 | 2F4U | 2.02 | 2 | 0.66 | 0.67 | 0.69 |
| 2BE0 | 2O3X | 2.16 | 2 | 0.67 | 0.65 | 0.66 |
| 2BE0 | 3BNL | 3.15 | 2 | 0.84 | 0.86 | 0.86 |
| 2BE0 | 4F8U | 2.50 | 2 | 0.82 | 0.80 | 0.82 |
| 2BEE | 1O9M | 2.05 | 2 | 0.66 | 0.64 | 0.66 |
| 2BEE | 2ET8 | 2.21 | 2 | 0.69 | 0.63 | 0.66 |
| 2BEE | 2F4S | 2.32 | 2 | 0.71 | 0.70 | 0.70 |
| 2BEE | 2F4U | 2.02 | 2 | 0.66 | 0.66 | 0.68 |
| 2BEE | 2O3X | 2.16 | 2 | 0.67 | 0.66 | 0.65 |
| 2BEE | 3BNL | 3.15 | 2 | 0.84 | 0.85 | 0.86 |
| 2BEE | 4F8U | 2.49 | 2 | 0.91 | 0.89 | 0.91 |
| 2ESJ | 2F4S | 2.02 | 2 | 0.58 | 0.53 | 0.58 |
| 2ESJ | 3BNL | 2.83 | 2 | 0.78 | 0.81 | 0.82 |
| 2ESJ | 4F8U | 2.44 | 2 | 0.81 | 0.79 | 0.80 |
| 2ET3 | 1O9M | 2.31 | 2 | 0.71 | 0.71 | 0.72 |
| 2ET3 | 2ET8 | 2.35 | 2 | 0.70 | 0.67 | 0.69 |
| 2ET3 | 2F4S | 2.01 | 2 | 0.62 | 0.55 | 0.56 |
| 2ET3 | 2F4T | 2.13 | 2 | 0.65 | 0.59 | 0.61 |
| 2ET3 | 3BNL | 3.24 | 2 | 0.80 | 0.82 | 0.82 |
| 2ET3 | 4F8U | 2.50 | 2 | 0.92 | 0.88 | 0.89 |
| 2ET4 | 3BNL | 2.88 | 2 | 0.74 | 0.78 | 0.79 |
| 2ET4 | 4F8U | 2.11 | 2 | 0.79 | 0.79 | 0.79 |
| 2ET5 | 2ET8 | 2.05 | 2 | 0.60 | 0.54 | 0.60 |
| 2ET5 | 3BNL | 2.90 | 2 | 0.75 | 0.77 | 0.79 |
| 2ET5 | 4F8U | 2.46 | 2 | 0.85 | 0.83 | 0.85 |
| 2ET8 | 1J7T | 2.02 | 2 | 0.60 | 0.57 | 0.57 |
| 2ET8 | 1MWL | 2.24 | 2 | 0.66 | 0.64 | 0.64 |
| 2ET8 | 1YRJ | 2.44 | 2 | 0.63 | 0.61 | 0.62 |
| 2ET8 | 2BE0 | 2.23 | 2 | 0.69 | 0.66 | 0.66 |
| 2ET8 | 2BEE | 2.21 | 2 | 0.69 | 0.65 | 0.65 |
| 2ET8 | 2ET3 | 2.35 | 2 | 0.70 | 0.70 | 0.71 |
| 2ET8 | 2ET5 | 2.05 | 2 | 0.59 | 0.55 | 0.54 |
| 2ET8 | 2F4S | 2.53 | 2 | 0.36 | 0.44 | 0.45 |
| 2ET8 | 2F4T | 2.15 | 2 | 0.34 | 0.23 | 0.23 |
| 2ET8 | 2F4U | 2.57 | 2 | 0.43 | 0.51 | 0.51 |
| 2ET8 | 2O3X | 2.47 | 2 | 0.33 | 0.40 | 0.41 |
| 2ET8 | 2PWT | 2.25 | 2 | 0.72 | 0.70 | 0.71 |
| 2ET8 | 3BNL | 2.78 | 2 | 0.69 | 0.73 | 0.73 |
| 2ET8 | 4F8U | 2.61 | 2 | 0.69 | 0.65 | 0.66 |
| 2ET8 | 4F8V | 2.19 | 2 | 0.62 | 0.59 | 0.59 |
| 2F4S | 1O9M | 2.64 | 2 | 0.49 | 0.53 | 0.54 |
| 2F4S | 1YRJ | 2.51 | 2 | 0.60 | 0.62 | 0.62 |
| 2F4S | 2BE0 | 2.33 | 2 | 0.71 | 0.66 | 0.67 |
| 2F4S | 2BEE | 2.32 | 2 | 0.71 | 0.66 | 0.67 |

| Supplementary Table 4 (continued) | | | | | | |
|---|---|---|---|---|---|---|
| Input PDB code | Target PDB code | RMSD | Cluster index | ENCoM | Cut-ANM | PD-ANM |
| 2F4S | 2ESJ | 2.02 | 2 | 0.58 | 0.59 | 0.61 |
| 2F4S | 2ET3 | 2.01 | 2 | 0.63 | 0.60 | 0.60 |
| 2F4S | 2ET8 | 2.53 | 2 | 0.37 | 0.46 | 0.47 |
| 2F4S | 2F4T | 2.32 | 2 | 0.46 | 0.41 | 0.43 |
| 2F4S | 2PWT | 2.28 | 2 | 0.70 | 0.67 | 0.68 |
| 2F4S | 3BNL | 3.11 | 2 | 0.64 | 0.68 | 0.69 |
| 2F4S | 4F8U | 2.66 | 2 | 0.76 | 0.73 | 0.74 |
| 2F4T | 1O9M | 2.00 | 2 | 0.49 | 0.48 | 0.51 |
| 2F4T | 1YRJ | 2.47 | 2 | 0.50 | 0.47 | 0.48 |
| 2F4T | 2ET3 | 2.13 | 2 | 0.78 | 0.81 | 0.78 |
| 2F4T | 2ET8 | 2.15 | 2 | 0.49 | 0.44 | 0.46 |
| 2F4T | 2F4S | 2.32 | 2 | 0.52 | 0.59 | 0.60 |
| 2F4T | 2F4U | 2.35 | 2 | 0.56 | 0.61 | 0.61 |
| 2F4T | 2O3X | 2.23 | 2 | 0.48 | 0.55 | 0.57 |
| 2F4T | 3BNL | 3.00 | 2 | 0.60 | 0.65 | 0.64 |
| 2F4T | 4F8U | 2.39 | 2 | 0.79 | 0.76 | 0.79 |
| 2F4T | 4F8V | 2.17 | 2 | 0.79 | 0.73 | 0.73 |
| 2F4U | 1O9M | 2.65 | 2 | 0.50 | 0.57 | 0.56 |
| 2F4U | 1YRJ | 2.23 | 2 | 0.54 | 0.55 | 0.54 |
| 2F4U | 2BE0 | 2.02 | 2 | 0.65 | 0.65 | 0.68 |
| 2F4U | 2BEE | 2.02 | 2 | 0.65 | 0.65 | 0.67 |
| 2F4U | 2ET8 | 2.57 | 2 | 0.43 | 0.50 | 0.50 |
| 2F4U | 2F4T | 2.35 | 2 | 0.42 | 0.43 | 0.42 |
| 2F4U | 2PWT | 2.06 | 2 | 0.60 | 0.58 | 0.61 |
| 2F4U | 3BNL | 2.81 | 2 | 0.74 | 0.78 | 0.78 |
| 2F4U | 4F8U | 2.68 | 2 | 0.81 | 0.79 | 0.81 |
| 2O3X | 1O9M | 2.55 | 2 | 0.44 | 0.53 | 0.55 |
| 2O3X | 1YRJ | 2.38 | 2 | 0.57 | 0.62 | 0.62 |
| 2O3X | 2BE0 | 2.16 | 2 | 0.67 | 0.65 | 0.66 |
| 2O3X | 2BEE | 2.16 | 2 | 0.66 | 0.64 | 0.66 |
| 2O3X | 2ET8 | 2.47 | 2 | 0.34 | 0.46 | 0.47 |
| 2O3X | 2F4T | 2.23 | 2 | 0.37 | 0.37 | 0.39 |
| 2O3X | 2PWT | 2.13 | 2 | 0.66 | 0.66 | 0.68 |
| 2O3X | 3BNL | 3.12 | 2 | 0.66 | 0.69 | 0.69 |
| 2O3X | 4F8U | 2.57 | 2 | 0.77 | 0.77 | 0.78 |
| 2PWT | 1O9M | 2.18 | 2 | 0.71 | 0.72 | 0.73 |
| 2PWT | 1YRJ | 2.04 | 2 | 0.70 | 0.71 | 0.71 |
| 2PWT | 2ET8 | 2.25 | 2 | 0.71 | 0.65 | 0.68 |
| 2PWT | 2F4S | 2.28 | 2 | 0.69 | 0.72 | 0.72 |
| 2PWT | 2F4U | 2.06 | 2 | 0.60 | 0.60 | 0.62 |
| 2PWT | 2O3X | 2.13 | 2 | 0.65 | 0.69 | 0.68 |
| 2PWT | 3BNL | 3.17 | 2 | 0.80 | 0.81 | 0.82 |
| 2PWT | 4F8U | 2.68 | 2 | 0.93 | 0.91 | 0.91 |
| 3BNL | 1J7T | 2.90 | 2 | 0.78 | 0.68 | 0.72 |
| 3BNL | 1LC4 | 2.96 | 2 | 0.80 | 0.64 | 0.71 |
| 3BNL | 1MWL | 3.32 | 2 | 0.82 | 0.69 | 0.76 |
| 3BNL | 1O9M | 2.91 | 2 | 0.74 | 0.62 | 0.68 |
| 3BNL | 1YRJ | 3.44 | 2 | 0.78 | 0.73 | 0.75 |
| 3BNL | 2BE0 | 3.15 | 2 | 0.85 | 0.72 | 0.78 |
| 3BNL | 2BEE | 3.15 | 2 | 0.84 | 0.71 | 0.77 |

| Supplementary Table 4 (continued) | | | | | | |
|---|---|---|---|---|---|---|
| Input PDB code | Target PDB code | RMSD | Cluster index | ENCoM | Cut-ANM | PD-ANM |
| 3BNL | 2ESJ | 2.83 | 2 | 0.80 | 0.69 | 0.73 |
| 3BNL | 2ET3 | 3.24 | 2 | 0.81 | 0.74 | 0.77 |
| 3BNL | 2ET4 | 2.88 | 2 | 0.75 | 0.61 | 0.68 |
| 3BNL | 2ET5 | 2.90 | 2 | 0.76 | 0.63 | 0.68 |
| 3BNL | 2ET8 | 2.78 | 2 | 0.71 | 0.60 | 0.65 |
| 3BNL | 2F4S | 3.11 | 2 | 0.66 | 0.56 | 0.65 |
| 3BNL | 2F4T | 3.00 | 2 | 0.62 | 0.52 | 0.59 |
| 3BNL | 2F4U | 2.81 | 2 | 0.75 | 0.63 | 0.69 |
| 3BNL | 2O3X | 3.12 | 2 | 0.67 | 0.56 | 0.65 |
| 3BNL | 2PWT | 3.17 | 2 | 0.81 | 0.68 | 0.73 |
| 3BNL | 4F8U | 4.09 | 2 | 0.88 | 0.82 | 0.85 |
| 3BNL | 4F8V | 3.15 | 2 | 0.77 | 0.66 | 0.68 |
| 3BNL | 4P20 | 3.03 | 2 | 0.80 | 0.65 | 0.71 |
| 4F8U | 1J7T | 2.36 | 2 | 0.92 | 0.93 | 0.93 |
| 4F8U | 1LC4 | 2.07 | 2 | 0.89 | 0.91 | 0.92 |
| 4F8U | 1MWL | 2.25 | 2 | 0.90 | 0.92 | 0.92 |
| 4F8U | 1O9M | 2.59 | 2 | 0.73 | 0.75 | 0.74 |
| 4F8U | 1YRJ | 2.81 | 2 | 0.84 | 0.83 | 0.82 |
| 4F8U | 2BE0 | 2.50 | 2 | 0.89 | 0.91 | 0.91 |
| 4F8U | 2BEE | 2.49 | 2 | 0.89 | 0.91 | 0.91 |
| 4F8U | 2ESJ | 2.44 | 2 | 0.89 | 0.91 | 0.91 |
| 4F8U | 2ET3 | 2.50 | 2 | 0.94 | 0.94 | 0.94 |
| 4F8U | 2ET4 | 2.11 | 2 | 0.91 | 0.92 | 0.92 |
| 4F8U | 2ET5 | 2.46 | 2 | 0.92 | 0.94 | 0.94 |
| 4F8U | 2ET8 | 2.61 | 2 | 0.72 | 0.71 | 0.71 |
| 4F8U | 2F4S | 2.66 | 2 | 0.78 | 0.77 | 0.77 |
| 4F8U | 2F4T | 2.39 | 2 | 0.70 | 0.70 | 0.70 |
| 4F8U | 2F4U | 2.68 | 2 | 0.81 | 0.83 | 0.83 |
| 4F8U | 2O3X | 2.57 | 2 | 0.78 | 0.77 | 0.77 |
| 4F8U | 2PWT | 2.68 | 2 | 0.92 | 0.94 | 0.94 |
| 4F8U | 3BNL | 4.09 | 2 | 0.81 | 0.82 | 0.81 |
| 4F8U | 4F8V | 2.56 | 2 | 0.92 | 0.93 | 0.93 |
| 4F8U | 4P20 | 2.09 | 2 | 0.91 | 0.93 | 0.93 |
| 4F8V | 1O9M | 2.28 | 2 | 0.66 | 0.70 | 0.74 |
| 4F8V | 1YRJ | 2.02 | 2 | 0.83 | 0.82 | 0.84 |
| 4F8V | 2ET8 | 2.19 | 2 | 0.60 | 0.62 | 0.67 |
| 4F8V | 2F4T | 2.17 | 2 | 0.62 | 0.60 | 0.62 |
| 4F8V | 3BNL | 3.15 | 2 | 0.77 | 0.78 | 0.81 |
| 4F8V | 4F8U | 2.56 | 2 | 0.84 | 0.81 | 0.85 |
| 4P20 | 3BNL | 3.03 | 2 | 0.78 | 0.81 | 0.82 |
| 4P20 | 4F8U | 2.09 | 2 | 0.89 | 0.85 | 0.89 |
| 1NLC | 1XP7 | 23.52 | 3 | 0.80 | 0.66 | 0.69 |
| 1NLC | 1XPF | 23.96 | 3 | 0.79 | 0.65 | 0.69 |
| 1NLC | 1Y3S | 23.66 | 3 | 0.80 | 0.66 | 0.70 |
| 1NLC | 1YXP | 23.70 | 3 | 0.80 | 0.66 | 0.69 |
| 1NLC | 1ZCI | 23.34 | 3 | 0.80 | 0.65 | 0.69 |
| 1NLC | 2B8S | 23.41 | 3 | 0.80 | 0.66 | 0.70 |
| 1NLC | 2FCX | 22.97 | 3 | 0.81 | 0.68 | 0.72 |
| 1NLC | 2FCY | 23.08 | 3 | 0.81 | 0.67 | 0.71 |
| 1NLC | 2FCZ | 22.94 | 3 | 0.81 | 0.67 | 0.71 |

| Supplementary Table 4 (continued) | | | | | | |
|---|---|---|---|---|---|---|
| Input PDB code | Target PDB code | RMSD | Cluster index | ENCoM | Cut-ANM | PD-ANM |
| 1NLC | 2FD0 | 23.28 | 3 | 0.80 | 0.66 | 0.70 |
| 1NLC | 2QEK | 2.60 | 3 | 0.63 | 0.54 | 0.60 |
| 1NLC | 3C44 | 2.44 | 3 | 0.74 | 0.64 | 0.68 |
| 1NLC | 3DVV | 2.76 | 3 | 0.73 | 0.63 | 0.65 |
| 1O3Z | 1XP7 | 23.50 | 3 | 0.80 | 0.66 | 0.69 |
| 1O3Z | 1XPF | 23.93 | 3 | 0.79 | 0.65 | 0.68 |
| 1O3Z | 1Y3S | 23.64 | 3 | 0.80 | 0.66 | 0.69 |
| 1O3Z | 1YXP | 23.68 | 3 | 0.80 | 0.66 | 0.69 |
| 1O3Z | 1ZCI | 23.31 | 3 | 0.80 | 0.66 | 0.69 |
| 1O3Z | 2B8S | 23.38 | 3 | 0.80 | 0.66 | 0.69 |
| 1O3Z | 2FCX | 22.94 | 3 | 0.81 | 0.68 | 0.71 |
| 1O3Z | 2FCY | 23.06 | 3 | 0.81 | 0.67 | 0.71 |
| 1O3Z | 2FCZ | 22.91 | 3 | 0.81 | 0.67 | 0.71 |
| 1O3Z | 2FD0 | 23.25 | 3 | 0.80 | 0.66 | 0.70 |
| 1O3Z | 2QEK | 2.64 | 3 | 0.64 | 0.57 | 0.63 |
| 1O3Z | 3C44 | 2.33 | 3 | 0.72 | 0.62 | 0.65 |
| 1O3Z | 3DVV | 2.68 | 3 | 0.72 | 0.61 | 0.62 |
| 1XP7 | 1NLC | 23.52 | 3 | 0.79 | 0.65 | 0.71 |
| 1XP7 | 1O3Z | 23.50 | 3 | 0.79 | 0.65 | 0.71 |
| 1XP7 | 2QEK | 23.49 | 3 | 0.79 | 0.67 | 0.72 |
| 1XP7 | 3C44 | 23.02 | 3 | 0.78 | 0.66 | 0.72 |
| 1XP7 | 3DVV | 22.68 | 3 | 0.78 | 0.67 | 0.73 |
| 1XP7 | 462D | 23.54 | 3 | 0.79 | 0.65 | 0.71 |
| 1XPF | 1NLC | 23.96 | 3 | 0.80 | 0.67 | 0.71 |
| 1XPF | 1O3Z | 23.93 | 3 | 0.80 | 0.67 | 0.71 |
| 1XPF | 2QEK | 23.93 | 3 | 0.80 | 0.69 | 0.72 |
| 1XPF | 3C44 | 23.46 | 3 | 0.79 | 0.69 | 0.73 |
| 1XPF | 3DVV | 23.12 | 3 | 0.79 | 0.70 | 0.73 |
| 1XPF | 462D | 23.97 | 3 | 0.80 | 0.67 | 0.71 |
| 1Y3S | 1NLC | 23.66 | 3 | 0.79 | 0.66 | 0.72 |
| 1Y3S | 1O3Z | 23.64 | 3 | 0.79 | 0.66 | 0.72 |
| 1Y3S | 2QEK | 23.63 | 3 | 0.79 | 0.67 | 0.73 |
| 1Y3S | 3C44 | 23.16 | 3 | 0.78 | 0.67 | 0.73 |
| 1Y3S | 3DVV | 22.81 | 3 | 0.78 | 0.68 | 0.74 |
| 1Y3S | 462D | 23.67 | 3 | 0.79 | 0.66 | 0.72 |
| 1YXP | 1NLC | 23.70 | 3 | 0.79 | 0.66 | 0.72 |
| 1YXP | 1O3Z | 23.68 | 3 | 0.79 | 0.66 | 0.72 |
| 1YXP | 2QEK | 23.67 | 3 | 0.79 | 0.67 | 0.73 |
| 1YXP | 3C44 | 23.19 | 3 | 0.78 | 0.67 | 0.73 |
| 1YXP | 3DVV | 22.84 | 3 | 0.78 | 0.68 | 0.73 |
| 1YXP | 462D | 23.72 | 3 | 0.79 | 0.66 | 0.72 |
| 1ZCI | 1NLC | 23.34 | 3 | 0.83 | 0.61 | 0.71 |
| 1ZCI | 1O3Z | 23.31 | 3 | 0.83 | 0.61 | 0.71 |
| 1ZCI | 2FCX | 2.08 | 3 | 0.84 | 0.83 | 0.82 |
| 1ZCI | 2QEK | 23.36 | 3 | 0.82 | 0.62 | 0.72 |
| 1ZCI | 3C44 | 22.85 | 3 | 0.83 | 0.62 | 0.73 |
| 1ZCI | 3DVV | 22.52 | 3 | 0.83 | 0.63 | 0.73 |
| 1ZCI | 462D | 23.35 | 3 | 0.83 | 0.61 | 0.71 |
| 2B8S | 1NLC | 23.41 | 3 | 0.79 | 0.65 | 0.70 |
| 2B8S | 1O3Z | 23.38 | 3 | 0.79 | 0.65 | 0.70 |

| Supplementary Table 4 (continued) | | | | | | |
|---|---|---|---|---|---|---|
| Input PDB code | Target PDB code | RMSD | Cluster index | ENCoM | Cut-ANM | PD-ANM |
| 2B8S | 2QEK | 23.38 | 3 | 0.79 | 0.67 | 0.71 |
| 2B8S | 3C44 | 22.90 | 3 | 0.78 | 0.67 | 0.71 |
| 2B8S | 3DVV | 22.56 | 3 | 0.78 | 0.67 | 0.72 |
| 2B8S | 462D | 23.42 | 3 | 0.79 | 0.65 | 0.70 |
| 2FCX | 1NLC | 22.97 | 3 | 0.79 | 0.64 | 0.73 |
| 2FCX | 1O3Z | 22.94 | 3 | 0.79 | 0.64 | 0.73 |
| 2FCX | 1ZCI | 2.08 | 3 | 0.88 | 0.78 | 0.81 |
| 2FCX | 2QEK | 22.90 | 3 | 0.78 | 0.65 | 0.73 |
| 2FCX | 3C44 | 22.46 | 3 | 0.78 | 0.66 | 0.74 |
| 2FCX | 3DVV | 22.10 | 3 | 0.78 | 0.66 | 0.74 |
| 2FCX | 462D | 22.98 | 3 | 0.79 | 0.64 | 0.73 |
| 2FCY | 1NLC | 23.08 | 3 | 0.80 | 0.63 | 0.70 |
| 2FCY | 1O3Z | 23.06 | 3 | 0.79 | 0.63 | 0.69 |
| 2FCY | 2QEK | 23.03 | 3 | 0.80 | 0.64 | 0.70 |
| 2FCY | 3C44 | 22.55 | 3 | 0.79 | 0.64 | 0.70 |
| 2FCY | 3DVV | 22.22 | 3 | 0.79 | 0.65 | 0.71 |
| 2FCY | 462D | 23.10 | 3 | 0.80 | 0.63 | 0.69 |
| 2FCZ | 1NLC | 22.94 | 3 | 0.79 | 0.63 | 0.73 |
| 2FCZ | 1O3Z | 22.91 | 3 | 0.79 | 0.63 | 0.73 |
| 2FCZ | 2QEK | 22.91 | 3 | 0.79 | 0.64 | 0.73 |
| 2FCZ | 3C44 | 22.41 | 3 | 0.79 | 0.64 | 0.73 |
| 2FCZ | 3DVV | 22.06 | 3 | 0.79 | 0.65 | 0.73 |
| 2FCZ | 462D | 22.94 | 3 | 0.79 | 0.63 | 0.72 |
| 2FD0 | 1NLC | 23.28 | 3 | 0.80 | 0.65 | 0.71 |
| 2FD0 | 1O3Z | 23.25 | 3 | 0.80 | 0.65 | 0.71 |
| 2FD0 | 2QEK | 23.26 | 3 | 0.80 | 0.66 | 0.71 |
| 2FD0 | 3C44 | 22.76 | 3 | 0.80 | 0.66 | 0.72 |
| 2FD0 | 3DVV | 22.42 | 3 | 0.80 | 0.67 | 0.72 |
| 2FD0 | 462D | 23.28 | 3 | 0.80 | 0.65 | 0.71 |
| 2QEK | 1NLC | 2.60 | 3 | 0.67 | 0.61 | 0.59 |
| 2QEK | 1O3Z | 2.64 | 3 | 0.68 | 0.63 | 0.60 |
| 2QEK | 1XP7 | 23.49 | 3 | 0.80 | 0.64 | 0.69 |
| 2QEK | 1XPF | 23.93 | 3 | 0.79 | 0.64 | 0.69 |
| 2QEK | 1Y3S | 23.63 | 3 | 0.80 | 0.65 | 0.70 |
| 2QEK | 1YXP | 23.67 | 3 | 0.80 | 0.65 | 0.70 |
| 2QEK | 1ZCI | 23.36 | 3 | 0.80 | 0.64 | 0.69 |
| 2QEK | 2B8S | 23.38 | 3 | 0.80 | 0.65 | 0.70 |
| 2QEK | 2FCX | 22.90 | 3 | 0.81 | 0.67 | 0.72 |
| 2QEK | 2FCY | 23.03 | 3 | 0.80 | 0.66 | 0.71 |
| 2QEK | 2FCZ | 22.91 | 3 | 0.81 | 0.66 | 0.71 |
| 2QEK | 2FD0 | 23.26 | 3 | 0.80 | 0.65 | 0.70 |
| 2QEK | 3C44 | 3.19 | 3 | 0.79 | 0.78 | 0.79 |
| 2QEK | 3DVV | 3.17 | 3 | 0.79 | 0.78 | 0.79 |
| 2QEK | 462D | 2.93 | 3 | 0.75 | 0.69 | 0.67 |
| 3C44 | 1NLC | 2.44 | 3 | 0.70 | 0.59 | 0.61 |
| 3C44 | 1O3Z | 2.33 | 3 | 0.68 | 0.56 | 0.58 |
| 3C44 | 1XP7 | 23.02 | 3 | 0.80 | 0.62 | 0.64 |
| 3C44 | 1XPF | 23.46 | 3 | 0.79 | 0.61 | 0.63 |
| 3C44 | 1Y3S | 23.16 | 3 | 0.80 | 0.62 | 0.64 |
| 3C44 | 1YXP | 23.19 | 3 | 0.80 | 0.62 | 0.64 |

| Supplementary Table 4 (continued) | | | | | | |
|---|---|---|---|---|---|---|
| Input PDB code | Target PDB code | RMSD | Cluster index | ENCoM | Cut-ANM | PD-ANM |
| 3C44 | 1ZCI | 22.85 | 3 | 0.80 | 0.62 | 0.64 |
| 3C44 | 2B8S | 22.90 | 3 | 0.80 | 0.62 | 0.65 |
| 3C44 | 2FCX | 22.46 | 3 | 0.81 | 0.64 | 0.66 |
| 3C44 | 2FCY | 22.55 | 3 | 0.80 | 0.63 | 0.66 |
| 3C44 | 2FCZ | 22.41 | 3 | 0.81 | 0.64 | 0.66 |
| 3C44 | 2FD0 | 22.76 | 3 | 0.80 | 0.63 | 0.64 |
| 3C44 | 2QEK | 3.19 | 3 | 0.67 | 0.63 | 0.62 |
| 3C44 | 462D | 2.59 | 3 | 0.73 | 0.67 | 0.67 |
| 3DVV | 1NLC | 2.76 | 3 | 0.69 | 0.61 | 0.66 |
| 3DVV | 1O3Z | 2.68 | 3 | 0.68 | 0.59 | 0.65 |
| 3DVV | 1XP7 | 22.68 | 3 | 0.80 | 0.57 | 0.62 |
| 3DVV | 1XPF | 23.12 | 3 | 0.79 | 0.56 | 0.61 |
| 3DVV | 1Y3S | 22.81 | 3 | 0.81 | 0.57 | 0.62 |
| 3DVV | 1YXP | 22.84 | 3 | 0.81 | 0.57 | 0.62 |
| 3DVV | 1ZCI | 22.52 | 3 | 0.80 | 0.57 | 0.62 |
| 3DVV | 2B8S | 22.56 | 3 | 0.80 | 0.58 | 0.62 |
| 3DVV | 2FCX | 22.10 | 3 | 0.81 | 0.59 | 0.64 |
| 3DVV | 2FCY | 22.22 | 3 | 0.81 | 0.58 | 0.63 |
| 3DVV | 2FCZ | 22.06 | 3 | 0.81 | 0.59 | 0.64 |
| 3DVV | 2FD0 | 22.42 | 3 | 0.80 | 0.58 | 0.63 |
| 3DVV | 2QEK | 3.17 | 3 | 0.64 | 0.62 | 0.64 |
| 3DVV | 462D | 2.65 | 3 | 0.68 | 0.60 | 0.64 |
| 462D | 1XP7 | 23.54 | 3 | 0.80 | 0.66 | 0.69 |
| 462D | 1XPF | 23.97 | 3 | 0.79 | 0.66 | 0.68 |
| 462D | 1Y3S | 23.67 | 3 | 0.81 | 0.66 | 0.69 |
| 462D | 1YXP | 23.72 | 3 | 0.81 | 0.66 | 0.68 |
| 462D | 1ZCI | 23.35 | 3 | 0.80 | 0.66 | 0.69 |
| 462D | 2B8S | 23.42 | 3 | 0.80 | 0.67 | 0.69 |
| 462D | 2FCX | 22.98 | 3 | 0.81 | 0.68 | 0.70 |
| 462D | 2FCY | 23.10 | 3 | 0.81 | 0.67 | 0.70 |
| 462D | 2FCZ | 22.94 | 3 | 0.81 | 0.68 | 0.70 |
| 462D | 2FD0 | 23.28 | 3 | 0.80 | 0.67 | 0.69 |
| 462D | 2QEK | 2.93 | 3 | 0.72 | 0.63 | 0.69 |
| 462D | 3C44 | 2.59 | 3 | 0.77 | 0.72 | 0.72 |
| 462D | 3DVV | 2.65 | 3 | 0.72 | 0.60 | 0.62 |
| 1XPE | 2OIJ | 22.73 | 4 | 0.78 | 0.65 | 0.71 |
| 1XPE | 2OIY | 22.66 | 4 | 0.78 | 0.65 | 0.71 |
| 1XPE | 2OJ0 | 22.97 | 4 | 0.78 | 0.65 | 0.71 |
| 1XPE | 3FAR | 22.76 | 4 | 0.78 | 0.65 | 0.71 |
| 2B8R | 2OIJ | 22.79 | 4 | 0.79 | 0.66 | 0.71 |
| 2B8R | 2OIY | 22.72 | 4 | 0.79 | 0.66 | 0.71 |
| 2B8R | 2OJ0 | 23.03 | 4 | 0.78 | 0.66 | 0.71 |
| 2B8R | 3FAR | 22.82 | 4 | 0.79 | 0.66 | 0.71 |
| 2OIJ | 1XPE | 22.73 | 4 | 0.82 | 0.68 | 0.68 |
| 2OIJ | 2B8R | 22.79 | 4 | 0.82 | 0.68 | 0.68 |
| 2OIY | 1XPE | 22.66 | 4 | 0.82 | 0.66 | 0.68 |
| 2OIY | 2B8R | 22.72 | 4 | 0.82 | 0.66 | 0.68 |
| 2OJ0 | 1XPE | 22.97 | 4 | 0.82 | 0.68 | 0.70 |
| 2OJ0 | 2B8R | 23.03 | 4 | 0.82 | 0.68 | 0.70 |
| 3FAR | 1XPE | 22.76 | 4 | 0.82 | 0.67 | 0.70 |

| Supplementary Table 4 (continued) | | | | | | |
|---|---|---|---|---|---|---|
| Input PDB code | Target PDB code | RMSD | Cluster index | ENCoM | Cut-ANM | PD-ANM |
| 3FAR | 2B8R | 22.82 | 4 | 0.82 | 0.67 | 0.69 |
| 1ZX7 | 1ZZ5 | 2.15 | 5 | 0.87 | 0.81 | 0.82 |
| 1ZX7 | 2A04 | 2.17 | 5 | 0.86 | 0.81 | 0.83 |
| 1ZZ5 | 1ZX7 | 2.15 | 5 | 0.88 | 0.83 | 0.83 |
| 2A04 | 1ZX7 | 2.17 | 5 | 0.89 | 0.82 | 0.83 |
| 2FQN | 2G5K | 3.06 | 6 | 0.60 | 0.52 | 0.53 |
| 2FQN | 5XZ1 | 3.37 | 6 | 0.66 | 0.62 | 0.62 |
| 2G5K | 2FQN | 3.06 | 6 | 0.57 | 0.60 | 0.60 |
| 2G5K | 2O3W | 2.86 | 6 | 0.51 | 0.54 | 0.54 |
| 2G5K | 5XZ1 | 3.97 | 6 | 0.63 | 0.62 | 0.64 |
| 2O3W | 2G5K | 2.86 | 6 | 0.51 | 0.45 | 0.46 |
| 2O3W | 5XZ1 | 3.05 | 6 | 0.59 | 0.53 | 0.54 |
| 5XZ1 | 2FQN | 3.37 | 6 | 0.73 | 0.66 | 0.72 |
| 5XZ1 | 2G5K | 3.97 | 6 | 0.65 | 0.59 | 0.62 |
| 5XZ1 | 2O3W | 3.05 | 6 | 0.65 | 0.62 | 0.65 |
| 2GPM | 439D | 7.85 | 7 | 0.40 | 0.16 | 0.18 |
| 439D | 2GPM | 7.85 | 7 | 0.36 | 0.16 | 0.16 |
| 2NOK | 2PN4 | 2.02 | 8 | 0.88 | 0.83 | 0.87 |
| 2PN4 | 2NOK | 2.02 | 8 | 0.87 | 0.85 | 0.87 |
| 3BNQ | 3BNR | 5.86 | 9 | 0.74 | 0.66 | 0.68 |
| 3BNQ | 3BNS | 5.89 | 9 | 0.74 | 0.66 | 0.68 |
| 3BNR | 3BNQ | 5.86 | 9 | 0.73 | 0.67 | 0.69 |
| 3BNS | 3BNQ | 5.89 | 9 | 0.73 | 0.66 | 0.68 |
| 3CW5 | 5L4O | 2.89 | 10 | 0.87 | 0.79 | 0.82 |
| 3CW6 | 5L4O | 2.89 | 10 | 0.87 | 0.80 | 0.82 |
| 5L4O | 3CW5 | 2.89 | 10 | 0.81 | 0.78 | 0.78 |
| 5L4O | 3CW6 | 2.89 | 10 | 0.78 | 0.75 | 0.75 |
| 3GCA | 6VUH | 2.03 | 11 | 0.30 | 0.39 | 0.26 |
| 6VUH | 3GCA | 2.03 | 11 | 0.53 | 0.49 | 0.51 |
| 3L0U | 6Y3G | 2.52 | 12 | 0.73 | 0.69 | 0.73 |
| 6Y3G | 3L0U | 2.52 | 12 | 0.82 | 0.80 | 0.82 |
| 3OWI | 3OWZ | 3.04 | 13 | 0.81 | 0.76 | 0.77 |
| 3OWW | 3OWZ | 3.07 | 13 | 0.82 | 0.77 | 0.78 |
| 3OWZ | 3OWI | 3.04 | 13 | 0.84 | 0.77 | 0.81 |
| 3OWZ | 3OWW | 3.07 | 13 | 0.84 | 0.77 | 0.82 |
| 3OWZ | 3OX0 | 3.02 | 13 | 0.83 | 0.76 | 0.81 |
| 3OWZ | 3OXB | 3.06 | 13 | 0.83 | 0.76 | 0.81 |
| 3OWZ | 3OXD | 3.03 | 13 | 0.83 | 0.76 | 0.80 |
| 3OWZ | 3OXE | 3.01 | 13 | 0.84 | 0.77 | 0.81 |
| 3OWZ | 3OXJ | 3.01 | 13 | 0.84 | 0.77 | 0.81 |
| 3OWZ | 3OXM | 2.96 | 13 | 0.83 | 0.76 | 0.81 |
| 3OX0 | 3OWZ | 3.02 | 13 | 0.81 | 0.75 | 0.77 |
| 3OXB | 3OWZ | 3.06 | 13 | 0.81 | 0.75 | 0.77 |
| 3OXD | 3OWZ | 3.03 | 13 | 0.81 | 0.75 | 0.77 |
| 3OXE | 3OWZ | 3.01 | 13 | 0.81 | 0.76 | 0.77 |
| 3OXJ | 3OWZ | 3.01 | 13 | 0.81 | 0.76 | 0.77 |
| 3OXM | 3OWZ | 2.96 | 13 | 0.81 | 0.75 | 0.77 |
| 3TD0 | 3TD1 | 3.50 | 14 | 0.65 | 0.57 | 0.59 |
| 3TD1 | 3TD0 | 3.50 | 14 | 0.73 | 0.64 | 0.66 |
| 3WRU | 4PDQ | 3.34 | 15 | 0.77 | 0.77 | 0.78 |

| Supplementary Table 4 (continued) | | | | | | |
|---|---|---|---|---|---|---|
| Input PDB code | Target PDB code | RMSD | Cluster index | ENCoM | Cut-ANM | PD-ANM |
| 3WRU | 6JBG | 2.11 | 15 | 0.79 | 0.78 | 0.78 |
| 4GPY | 4PDQ | 3.78 | 15 | 0.70 | 0.68 | 0.69 |
| 4GPY | 6JBG | 2.77 | 15 | 0.75 | 0.77 | 0.76 |
| 4PDQ | 3WRU | 3.34 | 15 | 0.76 | 0.70 | 0.71 |
| 4PDQ | 4GPY | 3.78 | 15 | 0.71 | 0.62 | 0.64 |
| 4PDQ | 6JBG | 3.79 | 15 | 0.70 | 0.66 | 0.68 |
| 6JBG | 3WRU | 2.11 | 15 | 0.79 | 0.75 | 0.78 |
| 6JBG | 4GPY | 2.77 | 15 | 0.75 | 0.69 | 0.71 |
| 6JBG | 4PDQ | 3.79 | 15 | 0.73 | 0.69 | 0.71 |
| 4K31 | 4K32 | 3.18 | 16 | 0.75 | 0.73 | 0.74 |
| 4K32 | 4K31 | 3.18 | 16 | 0.75 | 0.73 | 0.75 |
| 4L81 | 4OQU | 2.22 | 17 | 0.84 | 0.80 | 0.81 |
| 4OQU | 4L81 | 2.22 | 17 | 0.85 | 0.78 | 0.79 |
| 4MSB | 6Z18 | 2.24 | 18 | 0.95 | 0.55 | 0.61 |
| 4MSR | 6WY3 | 2.10 | 18 | 0.90 | 0.53 | 0.61 |
| 4MSR | 6Z18 | 2.44 | 18 | 0.94 | 0.53 | 0.67 |
| 5TDK | 6Z18 | 2.38 | 18 | 0.92 | 0.50 | 0.71 |
| 6WY3 | 4MSR | 2.10 | 18 | 0.91 | 0.87 | 0.89 |
| 6Z18 | 4MSB | 2.24 | 18 | 0.95 | 0.94 | 0.94 |
| 6Z18 | 4MSR | 2.44 | 18 | 0.94 | 0.89 | 0.93 |
| 6Z18 | 5TDK | 2.38 | 18 | 0.91 | 0.89 | 0.91 |
| 4P3S | 4P3T | 3.40 | 19 | 0.68 | 0.63 | 0.64 |
| 4P3T | 4P3S | 3.40 | 19 | 0.68 | 0.60 | 0.66 |
| 4P3U | 4P43 | 2.70 | 20 | 0.86 | 0.83 | 0.83 |
| 4P43 | 4P3U | 2.70 | 20 | 0.89 | 0.86 | 0.90 |
| 4RZD | 6XKN | 2.13 | 21 | 0.81 | 0.71 | 0.71 |
| 6XKN | 4RZD | 2.13 | 21 | 0.81 | 0.70 | 0.71 |
| 6XKN | 6XKO | 2.07 | 21 | 0.82 | 0.72 | 0.72 |
| 6XKO | 6XKN | 2.07 | 21 | 0.80 | 0.72 | 0.72 |
| 4TZX | 5E54 | 3.68 | 22 | 0.71 | 0.56 | 0.56 |
| 4TZX | 5SWD | 3.70 | 22 | 0.71 | 0.55 | 0.56 |
| 4TZY | 5E54 | 3.74 | 22 | 0.71 | 0.58 | 0.58 |
| 4TZY | 5SWD | 3.75 | 22 | 0.72 | 0.57 | 0.58 |
| 4XNR | 5E54 | 3.73 | 22 | 0.73 | 0.57 | 0.57 |
| 4XNR | 5SWD | 3.74 | 22 | 0.73 | 0.56 | 0.57 |
| 5E54 | 4TZX | 3.68 | 22 | 0.57 | 0.49 | 0.49 |
| 5E54 | 4TZY | 3.74 | 22 | 0.59 | 0.51 | 0.51 |
| 5E54 | 4XNR | 3.73 | 22 | 0.42 | 0.36 | 0.36 |
| 5E54 | 5SWE | 3.75 | 22 | 0.58 | 0.50 | 0.50 |
| 5E54 | 5UZA | 3.78 | 22 | 0.59 | 0.52 | 0.52 |
| 5E54 | 6VWT | 3.78 | 22 | 0.49 | 0.43 | 0.43 |
| 5E54 | 6VWV | 4.02 | 22 | 0.61 | 0.55 | 0.55 |
| 5SWD | 4TZX | 3.70 | 22 | 0.57 | 0.49 | 0.49 |
| 5SWD | 4TZY | 3.75 | 22 | 0.58 | 0.50 | 0.50 |
| 5SWD | 4XNR | 3.74 | 22 | 0.42 | 0.36 | 0.36 |
| 5SWD | 5SWE | 3.76 | 22 | 0.58 | 0.50 | 0.50 |
| 5SWD | 5UZA | 3.78 | 22 | 0.59 | 0.51 | 0.51 |
| 5SWD | 6VWT | 3.79 | 22 | 0.49 | 0.43 | 0.43 |
| 5SWD | 6VWV | 4.04 | 22 | 0.61 | 0.55 | 0.55 |
| 5SWE | 5E54 | 3.75 | 22 | 0.75 | 0.58 | 0.58 |

| Supplementary Table 4 (continued) | | | | | | |
|---|---|---|---|---|---|---|
| Input PDB code | Target PDB code | RMSD | Cluster index | ENCoM | Cut-ANM | PD-ANM |
| 5SWE | 5SWD | 3.76 | 22 | 0.75 | 0.58 | 0.58 |
| 5UZA | 5E54 | 3.78 | 22 | 0.73 | 0.58 | 0.59 |
| 5UZA | 5SWD | 3.78 | 22 | 0.73 | 0.58 | 0.58 |
| 6VWT | 5E54 | 3.78 | 22 | 0.69 | 0.58 | 0.58 |
| 6VWT | 5SWD | 3.79 | 22 | 0.69 | 0.58 | 0.58 |
| 6VWV | 5E54 | 4.02 | 22 | 0.75 | 0.65 | 0.65 |
| 6VWV | 5SWD | 4.04 | 22 | 0.75 | 0.66 | 0.65 |
| 4Y1J | 6CB3 | 25.96 | 23 | 0.98 | 0.96 | 0.96 |
| 6CB3 | 4Y1J | 25.96 | 23 | 0.98 | 0.90 | 0.93 |
| 5ZEG | 5ZEI | 3.10 | 24 | 0.76 | 0.73 | 0.76 |
| 5ZEG | 5ZEJ | 3.19 | 24 | 0.74 | 0.70 | 0.72 |
| 5ZEG | 5ZEM | 3.01 | 24 | 0.77 | 0.76 | 0.77 |
| 5ZEI | 5ZEG | 3.10 | 24 | 0.77 | 0.80 | 0.80 |
| 5ZEJ | 5ZEG | 3.19 | 24 | 0.76 | 0.76 | 0.76 |
| 5ZEM | 5ZEG | 3.01 | 24 | 0.72 | 0.76 | 0.75 |
| 6C8D | 6CAB | 11.27 | 25 | 0.56 | 0.54 | 0.53 |
| 6CAB | 6C8D | 11.27 | 25 | 0.42 | 0.38 | 0.37 |
| 6DLR | 6DNR | 2.62 | 26 | 0.84 | 0.80 | 0.81 |
| 6DLS | 6DNR | 2.48 | 26 | 0.85 | 0.83 | 0.84 |
| 6DNR | 6DLR | 2.62 | 26 | 0.85 | 0.82 | 0.84 |
| 6DNR | 6DLS | 2.48 | 26 | 0.87 | 0.85 | 0.86 |
| 6E80 | 6E81 | 2.88 | 27 | 0.76 | 0.72 | 0.74 |
| 6E80 | 6E82 | 2.91 | 27 | 0.71 | 0.65 | 0.70 |
| 6E80 | 6E84 | 2.97 | 27 | 0.81 | 0.76 | 0.78 |
| 6E81 | 6E80 | 2.88 | 27 | 0.70 | 0.66 | 0.67 |
| 6E82 | 6E80 | 2.91 | 27 | 0.68 | 0.66 | 0.67 |
| 6E84 | 6E80 | 2.97 | 27 | 0.73 | 0.68 | 0.71 |
| 6N5K | 6N5O | 2.25 | 28 | 0.50 | 0.49 | 0.49 |
| 6N5O | 6N5K | 2.25 | 28 | 0.45 | 0.48 | 0.48 |
| 7EOI | 7EOK | 2.01 | 29 | 0.78 | 0.76 | 0.77 |
| 7EOK | 7EOI | 2.01 | 29 | 0.72 | 0.70 | 0.71 |

**Supplementary Table 5:** Performance on individual NMR ensembles for the ensemble variance benchmark.

| PDB code | Cluster index | ENCoM NCO | Cut-ANM NCO | PD-ANM NCO | ENCoM RMSIP | Cut-ANM RMSIP | PD-ANM RMSIP |
|---|---|---|---|---|---|---|---|
| 2LUB | 1 | 0.83 | 0.81 | 0.82 | 0.74 | 0.72 | 0.73 |
| 1F5G | 2 | 0.83 | 0.83 | 0.85 | 0.61 | 0.62 | 0.65 |
| 1F5H | 2 | 0.78 | 0.78 | 0.77 | 0.62 | 0.58 | 0.58 |
| 2KYD | 3 | 0.93 | 0.94 | 0.94 | 0.74 | 0.76 | 0.75 |
| 1AFX | 4 | 0.60 | 0.56 | 0.58 | 0.48 | 0.47 | 0.47 |
| 6U79 | 5 | 0.65 | 0.67 | 0.70 | 0.60 | 0.61 | 0.64 |
| 2M21 | 6 | 0.78 | 0.71 | 0.73 | 0.62 | 0.58 | 0.59 |
| 1QET | 7 | 0.75 | 0.88 | 0.80 | 0.61 | 0.69 | 0.69 |
| 2N7M | 8 | 0.87 | 0.85 | 0.86 | 0.76 | 0.74 | 0.75 |
| 5KQE | 9 | 0.57 | 0.51 | 0.51 | 0.59 | 0.53 | 0.55 |
| 2LUN | 10 | 0.92 | 0.91 | 0.91 | 0.85 | 0.85 | 0.84 |
| 6VZC | 10 | 0.94 | 0.92 | 0.92 | 0.81 | 0.79 | 0.79 |
| 2FDT | 11 | 0.68 | 0.68 | 0.72 | 0.63 | 0.62 | 0.66 |

| Supplementary Table 5 (continued) | | | | | | | |
|---|---|---|---|---|---|---|---|
| PDB code | Cluster index | ENCoM NCO | Cut-ANM NCO | PD-ANM NCO | ENCoM RMSIP | Cut-ANM RMSIP | PD-ANM RMSIP |
| 1JU1 | 12 | 0.95 | 0.91 | 0.92 | 0.82 | 0.76 | 0.77 |
| 6PK9 | 13 | 0.54 | 0.53 | 0.53 | 0.60 | 0.59 | 0.62 |
| 1K6G | 14 | 0.75 | 0.74 | 0.79 | 0.57 | 0.58 | 0.61 |
| 1B36 | 15 | 0.94 | 0.91 | 0.92 | 0.84 | 0.78 | 0.79 |
| 28SP | 16 | 0.75 | 0.75 | 0.75 | 0.71 | 0.72 | 0.72 |
| 1OW9 | 17 | 0.60 | 0.52 | 0.52 | 0.61 | 0.49 | 0.49 |
| 2JXQ | 18 | 0.57 | 0.45 | 0.49 | 0.49 | 0.46 | 0.48 |
| 2JXS | 18 | 0.83 | 0.87 | 0.85 | 0.63 | 0.64 | 0.64 |
| 6K84 | 19 | 0.32 | 0.36 | 0.37 | 0.37 | 0.41 | 0.40 |
| 2ADT | 20 | 0.90 | 0.88 | 0.88 | 0.75 | 0.70 | 0.71 |
| 2JYF | 20 | 0.90 | 0.84 | 0.86 | 0.80 | 0.73 | 0.74 |
| 2JYH | 20 | 0.65 | 0.44 | 0.48 | 0.75 | 0.55 | 0.58 |
| 2JYJ | 20 | 0.95 | 0.92 | 0.92 | 0.86 | 0.79 | 0.80 |
| 2RN1 | 21 | 0.75 | 0.70 | 0.71 | 0.69 | 0.67 | 0.70 |
| 1K4A | 22 | 0.80 | 0.80 | 0.80 | 0.55 | 0.53 | 0.54 |
| 6XWJ | 23 | 0.96 | 0.96 | 0.96 | 0.77 | 0.77 | 0.77 |
| 6XXA | 23 | 0.86 | 0.86 | 0.86 | 0.69 | 0.69 | 0.70 |
| 3PHP | 24 | 0.92 | 0.89 | 0.89 | 0.72 | 0.70 | 0.72 |
| 2MI0 | 25 | 0.91 | 0.87 | 0.88 | 0.69 | 0.61 | 0.63 |
| 1TBK | 26 | 0.84 | 0.85 | 0.85 | 0.65 | 0.63 | 0.64 |
| 1YN1 | 26 | 0.83 | 0.81 | 0.83 | 0.64 | 0.59 | 0.62 |
| 6N8F | 27 | 0.94 | 0.94 | 0.94 | 0.73 | 0.71 | 0.71 |
| 1GUC | 28 | 0.70 | 0.65 | 0.65 | 0.57 | 0.57 | 0.57 |
| 1BN0 | 29 | 0.81 | 0.80 | 0.81 | 0.62 | 0.63 | 0.63 |
| 1LMV | 30 | 0.92 | 0.92 | 0.92 | 0.73 | 0.71 | 0.69 |
| 1FHK | 31 | 0.69 | 0.65 | 0.59 | 0.57 | 0.55 | 0.52 |
| 2NBZ | 32 | 0.95 | 0.94 | 0.94 | 0.89 | 0.86 | 0.86 |
| 2GIO | 33 | 0.84 | 0.81 | 0.81 | 0.72 | 0.67 | 0.67 |
| 2GIP | 33 | 0.93 | 0.93 | 0.93 | 0.71 | 0.70 | 0.70 |
| 1SLP | 34 | 0.83 | 0.75 | 0.81 | 0.59 | 0.56 | 0.60 |
| 1A9L | 35 | 0.90 | 0.87 | 0.88 | 0.78 | 0.74 | 0.75 |
| 1U3K | 35 | 0.92 | 0.90 | 0.90 | 0.79 | 0.74 | 0.74 |
| 2LX1 | 36 | 0.82 | 0.78 | 0.83 | 0.67 | 0.63 | 0.66 |
| 2KVN | 37 | 0.80 | 0.79 | 0.79 | 0.53 | 0.52 | 0.51 |
| 6BG9 | 38 | 0.99 | 0.95 | 0.95 | 0.99 | 0.95 | 0.95 |
| 2Y95 | 39 | 0.75 | 0.71 | 0.72 | 0.64 | 0.59 | 0.61 |
| 2LP9 | 40 | 0.64 | 0.56 | 0.59 | 0.61 | 0.53 | 0.54 |
| 2LPA | 40 | 0.71 | 0.71 | 0.72 | 0.60 | 0.63 | 0.63 |
| 1HLX | 41 | 0.90 | 0.83 | 0.82 | 0.65 | 0.58 | 0.58 |
| 1E4P | 42 | 0.94 | 0.92 | 0.93 | 0.74 | 0.69 | 0.70 |
| 2K3Z | 43 | 0.58 | 0.55 | 0.57 | 0.53 | 0.51 | 0.52 |
| 2MXJ | 44 | 0.68 | 0.71 | 0.72 | 0.52 | 0.54 | 0.54 |
| 1EBQ | 45 | 0.80 | 0.69 | 0.75 | 0.75 | 0.64 | 0.72 |
| 1EBR | 45 | 0.84 | 0.78 | 0.80 | 0.75 | 0.66 | 0.68 |
| 1EBS | 45 | 0.84 | 0.76 | 0.79 | 0.79 | 0.67 | 0.70 |
| 5V17 | 46 | 0.64 | 0.58 | 0.59 | 0.67 | 0.61 | 0.61 |
| 6XWW | 47 | 0.72 | 0.71 | 0.70 | 0.71 | 0.67 | 0.68 |
| 6XXB | 47 | 0.70 | 0.69 | 0.69 | 0.64 | 0.62 | 0.63 |
| 1R2P | 48 | 0.95 | 0.91 | 0.92 | 0.89 | 0.84 | 0.85 |

| Supplementary Table 5 (continued) | | | | | | | |
|---|---|---|---|---|---|---|---|
| PDB code | Cluster index | ENCoM NCO | Cut-ANM NCO | PD-ANM NCO | ENCoM RMSIP | Cut-ANM RMSIP | PD-ANM RMSIP |
| 2LPS | 48 | 0.88 | 0.86 | 0.87 | 0.83 | 0.82 | 0.83 |
| 1N66 | 49 | 0.87 | 0.82 | 0.86 | 0.67 | 0.64 | 0.67 |
| 2GRW | 49 | 0.72 | 0.71 | 0.71 | 0.66 | 0.64 | 0.64 |
| 1ATW | 50 | 0.53 | 0.46 | 0.55 | 0.59 | 0.47 | 0.55 |
| 2L5Z | 51 | 0.75 | 0.69 | 0.71 | 0.63 | 0.59 | 0.59 |
| 1JU7 | 52 | 0.71 | 0.70 | 0.72 | 0.60 | 0.60 | 0.61 |
| 2M23 | 53 | 0.96 | 0.95 | 0.96 | 0.83 | 0.82 | 0.82 |
| 2L8F | 54 | 0.87 | 0.85 | 0.85 | 0.76 | 0.72 | 0.73 |
| 1IDV | 55 | 0.55 | 0.30 | 0.37 | 0.47 | 0.41 | 0.40 |
| 2LBL | 56 | 0.65 | 0.65 | 0.69 | 0.58 | 0.63 | 0.65 |
| 2N3Q | 57 | 0.96 | 0.94 | 0.95 | 0.86 | 0.82 | 0.84 |
| 2M24 | 58 | 0.77 | 0.70 | 0.74 | 0.61 | 0.54 | 0.58 |
| 2IXY | 59 | 0.75 | 0.70 | 0.70 | 0.62 | 0.58 | 0.57 |
| 2N6X | 60 | 0.96 | 0.94 | 0.95 | 0.91 | 0.84 | 0.90 |
| 2IRO | 61 | 0.56 | 0.54 | 0.54 | 0.52 | 0.50 | 0.51 |
| 1Z31 | 62 | 0.66 | 0.56 | 0.57 | 0.64 | 0.56 | 0.57 |
| 2NC1 | 63 | 0.96 | 0.93 | 0.94 | 0.92 | 0.89 | 0.89 |
| 4A4S | 64 | 0.90 | 0.86 | 0.87 | 0.82 | 0.77 | 0.77 |
| 4A4T | 64 | 0.88 | 0.86 | 0.86 | 0.79 | 0.76 | 0.77 |
| 4A4U | 64 | 0.93 | 0.93 | 0.92 | 0.81 | 0.79 | 0.79 |
| 2N4L | 65 | 0.69 | 0.66 | 0.67 | 0.63 | 0.60 | 0.61 |
| 2F88 | 66 | 0.78 | 0.76 | 0.77 | 0.75 | 0.73 | 0.74 |
| 2LPT | 66 | 0.91 | 0.89 | 0.90 | 0.80 | 0.79 | 0.79 |
| 2LC8 | 67 | 0.74 | 0.60 | 0.61 | 0.65 | 0.56 | 0.57 |
| 2NCI | 68 | 0.77 | 0.70 | 0.72 | 0.72 | 0.64 | 0.65 |
| 1YLG | 69 | 0.63 | 0.56 | 0.56 | 0.58 | 0.54 | 0.54 |
| 1YNC | 69 | 0.71 | 0.66 | 0.67 | 0.58 | 0.54 | 0.56 |
| 1YNE | 69 | 0.67 | 0.64 | 0.66 | 0.56 | 0.51 | 0.53 |
| 1YNG | 69 | 0.59 | 0.58 | 0.58 | 0.55 | 0.55 | 0.56 |
| 1TJZ | 70 | 0.91 | 0.88 | 0.90 | 0.63 | 0.61 | 0.63 |
| 1T4X | 71 | 0.64 | 0.38 | 0.40 | 0.66 | 0.47 | 0.49 |
| 1MFJ | 72 | 0.76 | 0.78 | 0.78 | 0.65 | 0.66 | 0.67 |
| 2D18 | 73 | 0.90 | 0.92 | 0.93 | 0.72 | 0.75 | 0.78 |
| 2D19 | 73 | 0.86 | 0.83 | 0.84 | 0.73 | 0.68 | 0.69 |
| 2RLU | 74 | 0.56 | 0.55 | 0.58 | 0.57 | 0.56 | 0.57 |
| 1F7F | 75 | 0.90 | 0.86 | 0.87 | 0.78 | 0.72 | 0.73 |
| 1F7G | 75 | 0.88 | 0.81 | 0.83 | 0.71 | 0.62 | 0.64 |
| 6HYK | 76 | 0.78 | 0.81 | 0.87 | 0.78 | 0.76 | 0.78 |
| 2AHT | 77 | 0.94 | 0.77 | 0.86 | 0.76 | 0.63 | 0.68 |
| 2M22 | 78 | 0.76 | 0.75 | 0.76 | 0.60 | 0.58 | 0.60 |
| 1RNG | 79 | 0.53 | 0.53 | 0.51 | 0.52 | 0.51 | 0.51 |
| 1ATO | 80 | 0.65 | 0.66 | 0.65 | 0.53 | 0.53 | 0.53 |
| 2KHY | 81 | 0.93 | 0.91 | 0.91 | 0.76 | 0.72 | 0.72 |
| 1ZIF | 82 | 0.56 | 0.55 | 0.57 | 0.53 | 0.50 | 0.51 |
| 1WKS | 83 | 0.61 | 0.67 | 0.68 | 0.45 | 0.51 | 0.53 |
| 1XHP | 84 | 0.78 | 0.74 | 0.74 | 0.70 | 0.64 | 0.64 |
| 2O33 | 85 | 0.69 | 0.70 | 0.70 | 0.56 | 0.56 | 0.55 |
| 1A60 | 86 | 0.89 | 0.87 | 0.89 | 0.72 | 0.68 | 0.70 |
| 2GM0 | 87 | 0.91 | 0.89 | 0.89 | 0.84 | 0.81 | 0.82 |

| Supplementary Table 5 (continued) | | | | | | | |
|---|---|---|---|---|---|---|---|
| PDB code | Cluster index | ENCoM NCO | Cut-ANM NCO | PD-ANM NCO | ENCoM RMSIP | Cut-ANM RMSIP | PD-ANM RMSIP |
| 1Q75 | 88 | 0.60 | 0.70 | 0.70 | 0.50 | 0.52 | 0.53 |
| 2LV0 | 89 | 0.93 | 0.93 | 0.92 | 0.77 | 0.79 | 0.78 |
| 1F85 | 90 | 0.85 | 0.85 | 0.85 | 0.61 | 0.61 | 0.59 |
| 5LSN | 91 | 0.91 | 0.91 | 0.91 | 0.90 | 0.90 | 0.90 |
| 2M4W | 92 | 0.62 | 0.52 | 0.53 | 0.55 | 0.52 | 0.52 |
| 1ZIG | 93 | 0.71 | 0.71 | 0.72 | 0.58 | 0.58 | 0.60 |
| 2N6S | 94 | 0.91 | 0.87 | 0.86 | 0.83 | 0.81 | 0.80 |
| 6GE1 | 95 | 0.57 | 0.61 | 0.57 | 0.53 | 0.53 | 0.52 |
| 5A17 | 96 | 0.60 | 0.58 | 0.59 | 0.58 | 0.56 | 0.58 |
| 5A18 | 96 | 0.62 | 0.60 | 0.64 | 0.64 | 0.62 | 0.66 |
| 1LDZ | 97 | 0.90 | 0.84 | 0.87 | 0.65 | 0.59 | 0.61 |
| 2KY1 | 98 | 0.81 | 0.79 | 0.78 | 0.77 | 0.74 | 0.73 |
| 1IE1 | 99 | 0.55 | 0.52 | 0.52 | 0.48 | 0.43 | 0.43 |
| 1IE2 | 99 | 0.64 | 0.64 | 0.65 | 0.51 | 0.50 | 0.51 |
| 2MXL | 100 | 0.85 | 0.84 | 0.84 | 0.71 | 0.70 | 0.70 |
| 2FEY | 101 | 0.75 | 0.69 | 0.71 | 0.66 | 0.63 | 0.64 |
| 1F84 | 102 | 0.84 | 0.85 | 0.86 | 0.65 | 0.64 | 0.66 |
| 1QES | 103 | 0.88 | 0.86 | 0.86 | 0.69 | 0.69 | 0.69 |
| 2G1W | 104 | 0.66 | 0.64 | 0.66 | 0.55 | 0.56 | 0.56 |
| 1TXS | 105 | 0.76 | 0.70 | 0.72 | 0.64 | 0.59 | 0.61 |
| 1JO7 | 106 | 0.90 | 0.84 | 0.86 | 0.69 | 0.60 | 0.63 |
| 1MFY | 106 | 0.88 | 0.82 | 0.83 | 0.75 | 0.66 | 0.68 |
| 5VH7 | 107 | 0.93 | 0.92 | 0.92 | 0.74 | 0.74 | 0.73 |
| 2KOC | 108 | 0.68 | 0.71 | 0.71 | 0.54 | 0.55 | 0.54 |
| 6BY4 | 108 | 0.63 | 0.74 | 0.69 | 0.53 | 0.61 | 0.57 |
| 6BY5 | 108 | 0.55 | 0.61 | 0.59 | 0.51 | 0.55 | 0.52 |
| 5V2R | 109 | 0.93 | 0.91 | 0.91 | 0.68 | 0.63 | 0.64 |
| 2EUY | 110 | 0.92 | 0.91 | 0.92 | 0.75 | 0.72 | 0.72 |
| 1AQO | 111 | 0.83 | 0.75 | 0.77 | 0.65 | 0.57 | 0.59 |
| 1NBR | 111 | 0.72 | 0.68 | 0.68 | 0.64 | 0.59 | 0.60 |
| 1VOP | 112 | 0.31 | 0.42 | 0.49 | 0.41 | 0.49 | 0.55 |
| 1QWA | 113 | 0.86 | 0.83 | 0.83 | 0.67 | 0.66 | 0.66 |
| 2HNS | 114 | 0.92 | 0.89 | 0.91 | 0.70 | 0.66 | 0.68 |
| 1A3M | 115 | 0.79 | 0.81 | 0.81 | 0.69 | 0.73 | 0.74 |
| 2NBX | 116 | 0.92 | 0.89 | 0.89 | 0.89 | 0.85 | 0.86 |
| 5N5C | 117 | 0.77 | 0.74 | 0.75 | 0.50 | 0.48 | 0.48 |
| 1R7W | 118 | 0.82 | 0.76 | 0.77 | 0.65 | 0.57 | 0.58 |
| 1R7Z | 118 | 0.89 | 0.85 | 0.85 | 0.76 | 0.67 | 0.69 |
| 2NC0 | 119 | 0.93 | 0.91 | 0.92 | 0.79 | 0.75 | 0.76 |
| 6N8I | 120 | 0.93 | 0.91 | 0.91 | 0.74 | 0.71 | 0.71 |
| 2KPC | 121 | 0.76 | 0.73 | 0.73 | 0.55 | 0.51 | 0.53 |
| 2JXV | 122 | 0.88 | 0.89 | 0.89 | 0.80 | 0.80 | 0.81 |
| 2KRQ | 123 | 0.73 | 0.65 | 0.67 | 0.63 | 0.58 | 0.61 |
| 1M82 | 124 | 0.86 | 0.85 | 0.85 | 0.60 | 0.60 | 0.60 |
| 2MFD | 125 | 0.64 | 0.65 | 0.65 | 0.53 | 0.54 | 0.53 |
| 1P5O | 126 | 0.93 | 0.90 | 0.91 | 0.85 | 0.80 | 0.82 |
| 5KMZ | 127 | 0.60 | 0.60 | 0.61 | 0.58 | 0.56 | 0.58 |
| 2M18 | 128 | 0.53 | 0.55 | 0.57 | 0.49 | 0.55 | 0.56 |
| 6AAS | 129 | 0.64 | 0.57 | 0.65 | 0.46 | 0.49 | 0.51 |

| Supplementary Table 5 (continued) | | | | | | | |
|---|---|---|---|---|---|---|---|
| PDB code | Cluster index | ENCoM NCO | Cut-ANM NCO | PD-ANM NCO | ENCoM RMSIP | Cut-ANM RMSIP | PD-ANM RMSIP |
| 2L1F | 130 | 0.62 | 0.60 | 0.60 | 0.58 | 0.57 | 0.58 |
| 2RQJ | 131 | 0.39 | 0.34 | 0.35 | 0.45 | 0.41 | 0.40 |
| 2PCV | 132 | 0.93 | 0.91 | 0.91 | 0.82 | 0.80 | 0.80 |
| 2LK3 | 133 | 0.81 | 0.73 | 0.83 | 0.75 | 0.72 | 0.76 |
| 1YSV | 134 | 0.70 | 0.65 | 0.69 | 0.62 | 0.56 | 0.59 |
| 2MXK | 135 | 0.75 | 0.75 | 0.74 | 0.59 | 0.60 | 0.60 |
| 1MT4 | 136 | 0.83 | 0.80 | 0.81 | 0.61 | 0.58 | 0.61 |
| 2LJJ | 137 | 0.65 | 0.63 | 0.64 | 0.61 | 0.58 | 0.59 |
| 1P5M | 138 | 0.93 | 0.88 | 0.89 | 0.81 | 0.74 | 0.75 |
| 2RVO | 139 | 0.65 | 0.61 | 0.63 | 0.60 | 0.57 | 0.60 |
| 2IXZ | 140 | 0.39 | 0.47 | 0.52 | 0.38 | 0.47 | 0.47 |
| 1LC6 | 141 | 0.75 | 0.74 | 0.75 | 0.59 | 0.58 | 0.59 |
| 1NC0 | 141 | 0.74 | 0.70 | 0.71 | 0.62 | 0.58 | 0.58 |
| 1SY4 | 141 | 0.86 | 0.85 | 0.85 | 0.69 | 0.66 | 0.66 |
| 1SYZ | 141 | 0.63 | 0.65 | 0.66 | 0.62 | 0.59 | 0.60 |
| 2KEZ | 141 | 0.73 | 0.68 | 0.69 | 0.66 | 0.61 | 0.61 |
| 2KF0 | 141 | 0.77 | 0.70 | 0.74 | 0.76 | 0.71 | 0.72 |
| 1M5L | 142 | 0.95 | 0.89 | 0.89 | 0.83 | 0.77 | 0.77 |
| 1YMO | 143 | 0.79 | 0.75 | 0.77 | 0.65 | 0.60 | 0.62 |
| 2K95 | 143 | 0.83 | 0.80 | 0.80 | 0.66 | 0.63 | 0.63 |
| 2K96 | 143 | 0.76 | 0.75 | 0.75 | 0.64 | 0.63 | 0.63 |
| 7DD4 | 144 | 0.74 | 0.72 | 0.75 | 0.70 | 0.69 | 0.72 |
| 2MNC | 145 | 0.84 | 0.81 | 0.82 | 0.57 | 0.51 | 0.54 |
| 2LQZ | 146 | 0.69 | 0.61 | 0.65 | 0.61 | 0.55 | 0.58 |
| 1S34 | 147 | 0.44 | 0.35 | 0.35 | 0.49 | 0.49 | 0.49 |
| 1N8X | 148 | 0.85 | 0.84 | 0.85 | 0.74 | 0.71 | 0.73 |
| 2KUW | 149 | 0.95 | 0.90 | 0.90 | 0.87 | 0.79 | 0.80 |
| 2K5Z | 150 | 0.74 | 0.76 | 0.77 | 0.69 | 0.70 | 0.71 |
| 2OJ7 | 151 | 0.24 | 0.45 | 0.42 | 0.37 | 0.40 | 0.39 |
| 2OJ8 | 151 | 0.42 | 0.49 | 0.53 | 0.40 | 0.48 | 0.49 |
| 2LDL | 152 | 0.84 | 0.72 | 0.77 | 0.74 | 0.65 | 0.69 |
| 2MTJ | 153 | 0.97 | 0.94 | 0.95 | 0.91 | 0.85 | 0.86 |
| 5IEM | 154 | 0.89 | 0.85 | 0.87 | 0.82 | 0.78 | 0.80 |
| 1P5N | 155 | 0.77 | 0.74 | 0.77 | 0.64 | 0.61 | 0.64 |
| 5VH8 | 156 | 0.91 | 0.90 | 0.90 | 0.71 | 0.72 | 0.71 |
| 2D17 | 157 | 0.81 | 0.74 | 0.76 | 0.69 | 0.63 | 0.65 |
| 2JSE | 158 | 0.70 | 0.71 | 0.76 | 0.67 | 0.68 | 0.71 |
| 2KZL | 159 | 0.89 | 0.87 | 0.87 | 0.82 | 0.81 | 0.80 |
| 1CQL | 160 | 0.92 | 0.88 | 0.89 | 0.80 | 0.76 | 0.77 |
| 6VAR | 161 | 0.87 | 0.86 | 0.87 | 0.78 | 0.77 | 0.78 |
| 2PCW | 162 | 0.90 | 0.89 | 0.86 | 0.74 | 0.72 | 0.68 |
| 2KUU | 163 | 0.95 | 0.93 | 0.94 | 0.89 | 0.85 | 0.85 |
| 6W3M | 164 | 0.72 | 0.71 | 0.72 | 0.70 | 0.68 | 0.71 |
| 1OSW | 165 | 0.69 | 0.69 | 0.68 | 0.56 | 0.54 | 0.53 |
| 1SCL | 166 | 0.93 | 0.88 | 0.87 | 0.83 | 0.80 | 0.79 |
| 2IRN | 167 | 0.60 | 0.62 | 0.59 | 0.56 | 0.59 | 0.59 |
| 2KE6 | 168 | 0.96 | 0.95 | 0.95 | 0.90 | 0.88 | 0.88 |
| 2KUR | 168 | 0.94 | 0.93 | 0.93 | 0.84 | 0.83 | 0.83 |
| 1Z2J | 169 | 0.94 | 0.93 | 0.93 | 0.81 | 0.78 | 0.79 |

| Supplementary Table 5 (continued) | | | | | | | |
|---|---|---|---|---|---|---|---|
| PDB code | Cluster index | ENCoM NCO | Cut-ANM NCO | PD-ANM NCO | ENCoM RMSIP | Cut-ANM RMSIP | PD-ANM RMSIP |
| 2GV4 | 170 | 0.82 | 0.70 | 0.81 | 0.72 | 0.68 | 0.73 |
| 1K4B | 171 | 0.45 | 0.41 | 0.44 | 0.54 | 0.50 | 0.52 |
| 1BVJ | 172 | 0.70 | 0.62 | 0.68 | 0.62 | 0.57 | 0.63 |
| 2KD8 | 173 | 0.78 | 0.74 | 0.75 | 0.66 | 0.62 | 0.64 |
| 2P89 | 174 | 0.91 | 0.87 | 0.89 | 0.77 | 0.71 | 0.73 |
| 1K6H | 175 | 0.79 | 0.79 | 0.79 | 0.58 | 0.61 | 0.60 |
| 2KRP | 176 | 0.86 | 0.68 | 0.75 | 0.65 | 0.57 | 0.58 |
| 6VA1 | 177 | 0.83 | 0.84 | 0.82 | 0.62 | 0.62 | 0.61 |
| 2JTP | 178 | 0.88 | 0.87 | 0.88 | 0.66 | 0.62 | 0.64 |
| 2N8V | 179 | 0.64 | 0.61 | 0.64 | 0.60 | 0.59 | 0.63 |
| 2MQT | 180 | 0.87 | 0.86 | 0.87 | 0.80 | 0.78 | 0.79 |
| 1QWB | 181 | 0.71 | 0.63 | 0.64 | 0.61 | 0.54 | 0.55 |
| 5UZT | 182 | 0.87 | 0.79 | 0.80 | 0.75 | 0.65 | 0.68 |
| 2L3E | 183 | 0.72 | 0.67 | 0.68 | 0.63 | 0.58 | 0.59 |
| 1ATV | 184 | 0.28 | 0.25 | 0.24 | 0.47 | 0.40 | 0.40 |
| 1IKD | 185 | 0.91 | 0.86 | 0.87 | 0.62 | 0.58 | 0.58 |
| 1I3X | 186 | 0.62 | 0.47 | 0.59 | 0.54 | 0.46 | 0.52 |
| 2GVO | 187 | 0.74 | 0.70 | 0.71 | 0.54 | 0.51 | 0.51 |
| 2JR4 | 188 | 0.64 | 0.61 | 0.59 | 0.55 | 0.50 | 0.49 |
| 1FYO | 189 | 0.93 | 0.93 | 0.93 | 0.72 | 0.73 | 0.72 |
| 1TLR | 190 | 0.69 | 0.65 | 0.67 | 0.58 | 0.53 | 0.57 |
| 2QH4 | 191 | 0.83 | 0.86 | 0.86 | 0.58 | 0.60 | 0.60 |
| 2LHP | 192 | 0.86 | 0.85 | 0.86 | 0.72 | 0.72 | 0.73 |
| 2K41 | 193 | 0.49 | 0.46 | 0.47 | 0.51 | 0.47 | 0.47 |
| 1MFK | 194 | 0.73 | 0.72 | 0.74 | 0.55 | 0.57 | 0.59 |
| 1KKA | 195 | 0.77 | 0.73 | 0.72 | 0.66 | 0.63 | 0.63 |
| 2H49 | 196 | 0.59 | 0.57 | 0.59 | 0.53 | 0.52 | 0.54 |
| 7K4L | 197 | 0.82 | 0.65 | 0.71 | 0.71 | 0.61 | 0.66 |
| 2MEQ | 198 | 0.65 | 0.59 | 0.63 | 0.57 | 0.53 | 0.55 |
| 2KPD | 199 | 0.70 | 0.68 | 0.69 | 0.57 | 0.54 | 0.55 |
| 5KH8 | 200 | 0.71 | 0.57 | 0.61 | 0.67 | 0.58 | 0.59 |
| 1OQ0 | 201 | 0.83 | 0.82 | 0.88 | 0.57 | 0.56 | 0.59 |
| 1NA2 | 202 | 0.92 | 0.92 | 0.92 | 0.77 | 0.74 | 0.74 |
| 2KXZ | 203 | 0.96 | 0.96 | 0.96 | 0.90 | 0.90 | 0.90 |
| 5WQ1 | 204 | 0.45 | 0.47 | 0.50 | 0.42 | 0.42 | 0.45 |
| 1ANR | 205 | 0.79 | 0.69 | 0.73 | 0.59 | 0.53 | 0.56 |
| 7JU1 | 205 | 0.76 | 0.67 | 0.69 | 0.56 | 0.49 | 0.50 |
| 2F87 | 206 | 0.54 | 0.55 | 0.54 | 0.49 | 0.51 | 0.51 |
| 2N6T | 207 | 0.86 | 0.84 | 0.84 | 0.79 | 0.76 | 0.76 |
| 2F4X | 208 | 0.84 | 0.90 | 0.91 | 0.64 | 0.66 | 0.66 |
| 1ELH | 209 | 0.91 | 0.78 | 0.86 | 0.84 | 0.74 | 0.79 |
| 1JOX | 210 | 0.75 | 0.66 | 0.69 | 0.56 | 0.55 | 0.54 |
| 1JP0 | 210 | 0.90 | 0.89 | 0.89 | 0.59 | 0.54 | 0.56 |
| 2JWV | 211 | 0.97 | 0.97 | 0.96 | 0.85 | 0.83 | 0.83 |
| 2L2J | 212 | 0.94 | 0.94 | 0.94 | 0.82 | 0.82 | 0.83 |
| 1RRR | 213 | 0.65 | 0.70 | 0.71 | 0.55 | 0.59 | 0.61 |
| 2K66 | 214 | 0.95 | 0.79 | 0.81 | 0.80 | 0.70 | 0.69 |
| 2M12 | 215 | 0.57 | 0.51 | 0.53 | 0.54 | 0.46 | 0.48 |
| 2N7X | 216 | 0.87 | 0.86 | 0.86 | 0.63 | 0.62 | 0.63 |

| Supplementary Table 5 (continued) | | | | | | | |
|---|---|---|---|---|---|---|---|
| PDB code | Cluster index | ENCoM NCO | Cut-ANM NCO | PD-ANM NCO | ENCoM RMSIP | Cut-ANM RMSIP | PD-ANM RMSIP |
| 1ESY | 217 | 0.59 | 0.60 | 0.63 | 0.50 | 0.52 | 0.54 |
| 1KKS | 218 | 0.79 | 0.57 | 0.56 | 0.66 | 0.52 | 0.52 |
| 2LU0 | 219 | 0.89 | 0.85 | 0.86 | 0.78 | 0.73 | 0.75 |
| 2KPV | 220 | 0.68 | 0.64 | 0.64 | 0.69 | 0.63 | 0.64 |
| 2K65 | 221 | 0.90 | 0.92 | 0.92 | 0.69 | 0.70 | 0.68 |
| 1QC8 | 222 | 0.72 | 0.64 | 0.66 | 0.51 | 0.44 | 0.46 |
| 5V16 | 223 | 0.77 | 0.73 | 0.72 | 0.81 | 0.78 | 0.79 |
| 2JYM | 224 | 0.85 | 0.83 | 0.84 | 0.68 | 0.64 | 0.67 |
| 1K5I | 225 | 0.83 | 0.82 | 0.83 | 0.72 | 0.70 | 0.71 |
| 2LAC | 226 | 0.86 | 0.82 | 0.86 | 0.70 | 0.63 | 0.70 |
| 1A51 | 227 | 0.97 | 0.94 | 0.94 | 0.90 | 0.84 | 0.86 |
| 1S9S | 228 | 0.94 | 0.90 | 0.91 | 0.89 | 0.85 | 0.86 |
| 1BGZ | 229 | 0.69 | 0.80 | 0.79 | 0.66 | 0.70 | 0.71 |
| 2XEB | 230 | 0.52 | 0.41 | 0.44 | 0.66 | 0.60 | 0.63 |
| 2KBP | 231 | 0.44 | 0.51 | 0.51 | 0.43 | 0.45 | 0.46 |
| 1ROQ | 232 | 0.85 | 0.88 | 0.88 | 0.65 | 0.61 | 0.61 |
| 2N6W | 233 | 0.94 | 0.93 | 0.93 | 0.86 | 0.82 | 0.83 |
| 2LBK | 234 | 0.66 | 0.65 | 0.68 | 0.59 | 0.61 | 0.61 |
| 2GV3 | 235 | 0.74 | 0.60 | 0.65 | 0.61 | 0.54 | 0.55 |
| 1ZC5 | 236 | 0.94 | 0.91 | 0.91 | 0.79 | 0.78 | 0.78 |
| 1MNX | 237 | 0.93 | 0.91 | 0.91 | 0.81 | 0.80 | 0.79 |
| 1JTJ | 238 | 0.91 | 0.87 | 0.88 | 0.64 | 0.56 | 0.59 |
| 1JUR | 239 | 0.70 | 0.65 | 0.66 | 0.65 | 0.63 | 0.63 |
| 1Z30 | 240 | 0.85 | 0.83 | 0.82 | 0.66 | 0.62 | 0.63 |
| 2M8K | 241 | 0.77 | 0.73 | 0.75 | 0.63 | 0.57 | 0.60 |
| 2M57 | 242 | 0.87 | 0.84 | 0.86 | 0.69 | 0.63 | 0.66 |
| 17RA | 243 | 0.97 | 0.96 | 0.96 | 0.82 | 0.80 | 0.79 |
| 2N2P | 244 | 0.74 | 0.54 | 0.60 | 0.64 | 0.51 | 0.54 |
| 2B7G | 245 | 0.65 | 0.60 | 0.64 | 0.66 | 0.61 | 0.63 |
| 1HWQ | 246 | 0.87 | 0.85 | 0.86 | 0.68 | 0.65 | 0.66 |
| 1ZIH | 247 | 0.64 | 0.62 | 0.63 | 0.56 | 0.53 | 0.53 |
| 1UUU | 248 | 0.69 | 0.58 | 0.63 | 0.56 | 0.50 | 0.55 |
| 1E95 | 249 | 0.89 | 0.82 | 0.85 | 0.71 | 0.64 | 0.66 |
| 2KRL | 250 | 0.70 | 0.64 | 0.65 | 0.69 | 0.63 | 0.64 |
| 2HEM | 251 | 0.87 | 0.79 | 0.81 | 0.74 | 0.70 | 0.72 |
| 2LDT | 252 | 0.81 | 0.78 | 0.80 | 0.68 | 0.63 | 0.64 |
| 2QH3 | 253 | 0.82 | 0.82 | 0.83 | 0.61 | 0.57 | 0.58 |
| 2RPT | 254 | 0.94 | 0.92 | 0.92 | 0.75 | 0.71 | 0.71 |
| 6MXQ | 255 | 0.88 | 0.86 | 0.86 | 0.82 | 0.81 | 0.81 |
| 5UF3 | 256 | 0.68 | 0.70 | 0.73 | 0.76 | 0.78 | 0.80 |
| 2L6I | 257 | 0.53 | 0.48 | 0.49 | 0.54 | 0.52 | 0.51 |
| 2HUA | 258 | 0.93 | 0.90 | 0.90 | 0.87 | 0.83 | 0.83 |
| 1D0U | 259 | 0.87 | 0.72 | 0.74 | 0.62 | 0.56 | 0.57 |
| 2M5U | 260 | 0.96 | 0.96 | 0.96 | 0.83 | 0.80 | 0.79 |
| 2EVY | 261 | 0.60 | 0.56 | 0.55 | 0.61 | 0.60 | 0.60 |
| 2QH2 | 262 | 0.88 | 0.83 | 0.85 | 0.65 | 0.61 | 0.62 |
| 2KY2 | 263 | 0.91 | 0.90 | 0.90 | 0.71 | 0.70 | 0.70 |
| 2N2O | 264 | 0.67 | 0.47 | 0.51 | 0.64 | 0.46 | 0.50 |
| 7LVA | 265 | 0.93 | 0.92 | 0.92 | 0.90 | 0.88 | 0.89 |

| Supplementary Table 5 (continued) | | | | | | | |
|---|---|---|---|---|---|---|---|
| PDB code | Cluster index | ENCoM NCO | Cut-ANM NCO | PD-ANM NCO | ENCoM RMSIP | Cut-ANM RMSIP | PD-ANM RMSIP |
| 6NOA | 266 | 0.83 | 0.82 | 0.83 | 0.76 | 0.74 | 0.76 |
| 2LBJ | 267 | 0.64 | 0.55 | 0.56 | 0.67 | 0.56 | 0.56 |
| 1PJY | 268 | 0.87 | 0.86 | 0.85 | 0.63 | 0.60 | 0.59 |
| 2ES5 | 269 | 0.81 | 0.62 | 0.66 | 0.64 | 0.52 | 0.54 |
| 2KY0 | 270 | 0.87 | 0.86 | 0.85 | 0.81 | 0.76 | 0.75 |
| 2NBY | 271 | 0.97 | 0.94 | 0.96 | 0.86 | 0.84 | 0.85 |

**Supplementary Table 6:** MD trajectories of miR-125a variants

| Base pair 22 | Replicate | Simulation time (ns) |
|---|---|---|
| AA | 1 | 122.6 |
| AA | 2 | 122.4 |
| AA | 3 | 120.3 |
| AC | 1 | 123.6 |
| AC | 2 | 135.4 |
| AC | 3 | 123.2 |
| AG | 1 | 107.4 |
| AG | 2 | 110.4 |
| AG | 3 | 121.2 |
| AU | 1 | 123.4 |
| AU | 2 | 123.2 |
| AU | 3 | 110.2 |
| CA | 1 | 123.2 |
| CA | 2 | 122.4 |
| CA | 3 | 123.3 |
| CC | 1 | 110.5 |
| CC | 2 | 111.1 |
| CC | 3 | 112.1 |
| CG | 1 | 135.4 |
| CG | 2 | 124.1 |
| CG | 3 | 123.2 |
| CU | 1 | 125.3 |
| CU | 2 | 111.4 |
| CU | 3 | 123.1 |
| GA | 1 | 109.5 |
| GA | 2 | 112.2 |
| GA | 3 | 109.1 |
| GC | 1 | 124.5 |
| GC | 2 | 124.1 |
| GC | 3 | 112.5 |
| GG | 1 | 112.4 |
| GG | 2 | 108.8 |
| GG | 3 | 137.4 |
| GU | 1 | 123.4 |
| GU | 2 | 111.5 |
| GU | 3 | 112.1 |
| UA | 1 | 112.4 |
| UA | 2 | 122.3 |

| Supplementary Table 6 (continued) | | |
|---|---|---|
| Base pair 22 | Replicate | Simulation time (ns) |
| UA | 3 | 122.2 |
| UC | 1 | 132.1 |
| UC | 2 | 134.1 |
| UC | 3 | 119.1 |
| UG | 1 | 112.1 |
| UG | 2 | 123.5 |
| UG | 3 | 124.3 |
| UU | 1 | 124.1 |
| UU | 2 | 121.2 |
| UU | 3 | 121.6 |

**Supplementary Table 7:** ENCoM interaction strength for different base pairs. The base pairs are taken from the WT miR-125a structure.

| Base pair | Interaction strength |
|---|---|
| AC | 134.9 |
| AG | 0.0 |
| AU | 167.8 |
| CG | 218.0 |
| CU | 150.7 |
| GU | 149.4 |
| UU | 53.7 |

[5] Paul Dallaire, Huiping Tan, Keith Szulwach, Christopher Ma, Peng Jin, and François Major. Structural dynamics control the MicroRNA maturation pathway. *Nucleic Acids Res*, page gkw793, sep 2016.
